# Neural trajectories reveal orchestration of cortical coding underlying natural language composition

**DOI:** 10.1101/2025.04.24.650382

**Authors:** Xiangrong Tang, James R. Booth, Yunhao Zhang, Shaonan Wang, Guosheng Ding

## Abstract

Language is a hierarchically structured system that enables humans to communicate complex meanings. Despite recent advances, the neurocomputational mechanism underlying the composition of natural language remains unclear. Building on the neural population theory, we investigated how neural trajectories in latent spaces underpin natural language composition that integrates diverse lexical content and syntactic relations. We found that neural trajectories derived from human neocortical responses show an orchestration of distinct coding strategies during naturalistic story comprehension. Neural latent geometry is primarily associated with syntactic relations and exhibits more efficient compression relative to lexical content. We further demonstrate that these trajectories can be simulated by brain-inspired computing systems with near-critical dynamics and a preference for historical information. Overall, by positioning structure-based integration as a key computation of natural language comprehension, our findings provide a novel perspective on the mechanism underlying real-world language use and emphasize the importance of contextual information in the development of brain-inspired intelligent systems.

## Introduction

Language allows humans to convey infinite meanings based on finite linguistic units and syntactic structures in natural contexts^1–4^. Successful understanding of language requires the continuous integration of lexical content and syntactic relations to accurately compose or reconstruct the conveyed meaning from ongoing speech streams. This composition of natural language is underpinned by complex neural orchestrations in the human brain, which require flexible, resilient and effective cortical coding to balance the constrained neural resources and the huge computational burden imposed by the infinite number of possible linguistic combinations. Previous imaging studies have identified an amodal, language-selective core network that accesses lexical information and integrates word meanings into more complex representations^5–9^. Other studies have revealed interleaved neural populations with different timescales within this language network^10,11^ and their dynamics of neural activity for language processing^12–16^. Notably, the language network interacts with widespread cortical regions in real-life language use, showing that neural substrates associated with lexical and syntactic information are distributed across the human neocortex^17–21^. Despite these advances, the cortical computation underlying the composition of natural language, which entails flexible integration of lexical content and syntactic relations, remains unclear.

Neural population theory provides a framework for understanding the computations of neural ensembles^22–24^ and offers us a new perspective for exploring the neural implementation of natural language composition. This theory posits that task-relevant information is represented by neural manifolds in low-dimensional latent spaces embedded in high-dimensional population activity, where neural computations are implemented as neural trajectories on these manifolds^25–27^. For example, hippocampal population activity forms a neural manifold that topologically mirrors the physical environment, with trajectories on this manifold reflecting the spatial navigation of rodents^28,29^. Benefiting from advances in latent variable modeling^30,31^, neural latent spaces provide a more interpretable scaffold for linking behavior to neural activity and uncovering neural computations. However, it remains poorly understood whether neural latent trajectories, primarily characterized in non-human animal models, can be generalized to explain cortical coding in humans that support higher-order functions including language composition. Here, we postulate that the composition of natural language is implemented by the formation of multidimensional latent representations, which are driven by the neural computation of integrating lexical and syntactic information. Under this framework, the composition of natural language can be studied by tracking neural trajectories, and the correspondence between specific linguistic information and cortical responses can be revealed by analyzing the geometry of latent spaces during the comprehension of naturalistic material.

In addition, the dimensionality of latent spaces provides an approach to unravel the neural coding strategies of natural language composition. We propose that the human neocortex may orchestrate distinct coding strategies to facilitate the construction of linguistic meaning. Here, we consider two classical strategies of population coding. The strategy of efficient coding decorrelates inputs and allows complex information in circuits to be decoded by downstream networks^32–34^, resulting in high-dimensional neural representations containing maximal information^35,36^. In contrast, the strategy of robust coding allows the circuits to compress a wide range of inputs into a minimal number of common components^37–39^. Neural responses with robust coding are correlated and redundant, resulting in low-dimensional and noise-resistant representations^40,41^. The dimensionality of neural representations has been used to study neural coding in various cognitive domains. Previous studies have shown that simple tasks with low cognitive demands are associated with low-dimensional neural patterns, whereas more complex and demanding tasks correspond to higher-dimensional patterns^16,42,43^. Based on these findings, we hypothesize that the human cortex may use robust coding to capture local lexical content and use efficient coding to enable the real-time establishment of syntactic dependency relations.

In this study, we investigated how the human neocortex performs natural language composition, which refers to the dynamic integration of lexical and syntactic information to construct diverse linguistic meanings, by examining neural trajectories in latent spaces. For this purpose, we used a recently published multimodal neuroimaging dataset containing magnetoencephalography (MEG) recordings from twelve participants^44^. During the MEG acquisition, they listened to 60 stories over six hours and naturally engaged in continuous combinatorial operations. First, we examined whether neural latent trajectories capture language-relevant information from naturalistic stories. Second, we tested the associations of lexical and syntactic features with neural latent geometry. If the composition of natural language primarily involves lexical content, there should be stronger relations between lexical patterns and neural latent patterns. Conversely, if syntactic relations are dominant, there should be stronger relations between syntactic patterns and neural latent patterns. We then investigated whether different coding strategies are used to integrate lexical content and syntactic relations. Namely, we compared the logarithmic model, associated with robust coding, with the first-order polynomial model, representing efficient coding, for lexical and syntactic patterns, respectively. Finally, we explored the computational principles underlying the human-like composition by investigating whether brain-inspired computing systems, which better store and integrate more long-range information, are more likely to generate observed neural trajectories.

## Results

### Neural trajectories in latent spaces capture consistent components across participants and separate individual words in naturalistic stories

To characterize neural trajectories in latent spaces of human neocortical responses during natural language comprehension, we applied a nonlinear encoder with time contrastive learning to parcellated MEG data based on a framework for estimating consistent embeddings of high-dimensional recordings using auxiliary variables (CEBRA; Fig. 1, Fig. 2a-d)^30^. We transformed these MEG data into neural trajectories as CEBRA-Time embeddings for each run, participant, and dimension setting (Fig. 3a). To obtain benchmarks, we computed low-dimensional embeddings using other dimensionality reduction techniques, including principal component analysis (PCA), independent component analysis (ICA), and uniform manifold approximation and projection (UMAP)^45^. We found that CEBRA-Time embeddings were more consistent across participants and better at discriminating individual words than those derived from other techniques.

**Figure 1.**
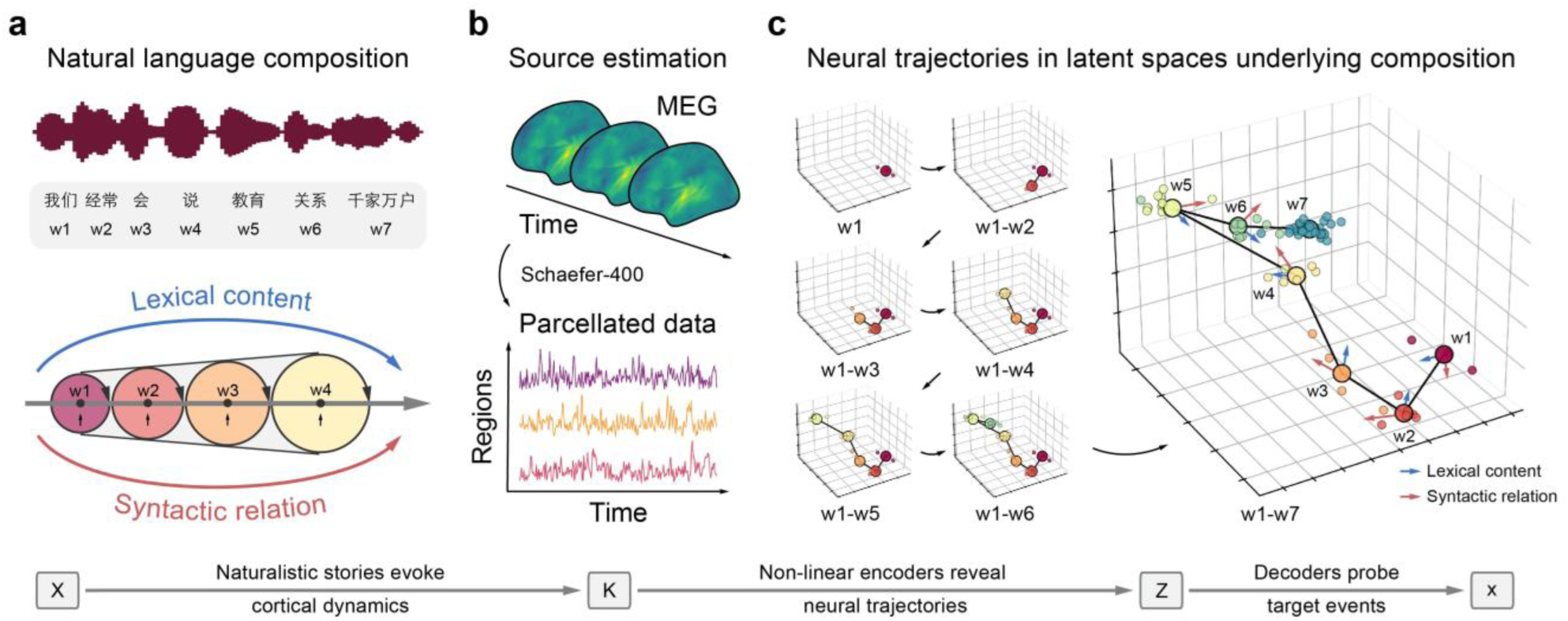
Overview of the procedure used to investigate neural trajectories in latent spaces underlying the composition of natural language. **a)** Twelve participants listened to 60 naturalistic stories (X) and naturally engaged in the integration of lexical content and syntactic relations. **b)** Human neocortical dynamics were observed as region-level MEG data (K) that were reconstructed from sensor-level data and parcellated into a predefined cortical atlas. **c)** Neural trajectories in latent spaces (Z) were transformed from region-level MEG data and were examined using decoders to reveal how specific linguistic features (x) were integrated into multidimensional latent representations. The neural trajectories visualized here are the group-averaged trajectories derived from the CEBRA-Behavior embeddings of the example sentence. Framed circles indicate the representations of individual words, while unframed circles indicate the neural states at each time point.

**Figure 2.**
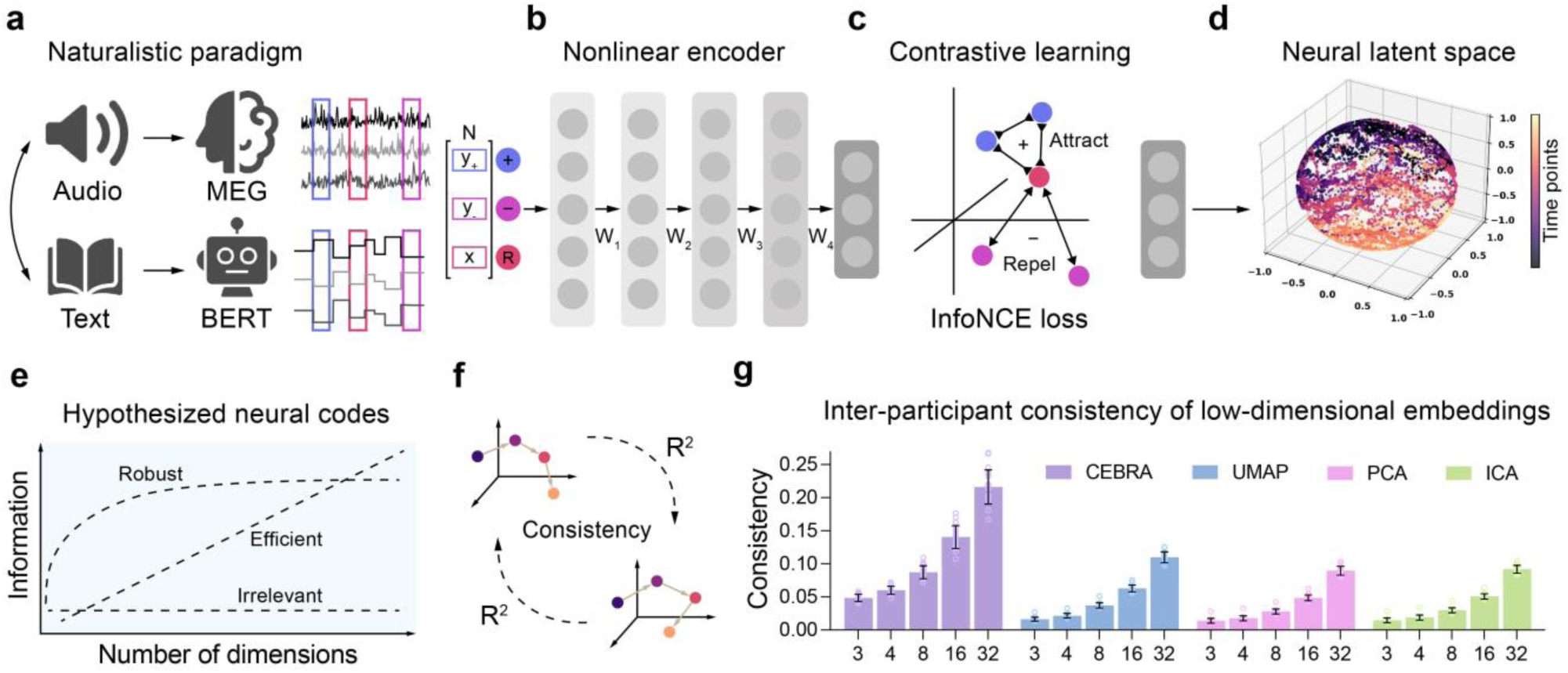
Generation of neural trajectories in the latent spaces from the region-level MEG data using the CEBRA framework. **a-d)** Nonlinear encoders with time/behavioral contrastive learning were used to transform region-level MEG data into neural trajectories in latent spaces for each participant listening to each naturalistic story (adapted from Schneider et al, 2024, under CC BY 4.0 license). **e)** Three relationships were evaluated to examine the cortical coding strategies that may facilitate natural language composition. **f)** Linear regression was performed between neural latent embeddings to assess their consistency across pairs of participants. **g)** The inter-participant consistency of neural embeddings derived from CEBRA-Time and other benchmarks. The *R*² values were averaged across all 132 directional pairs from twelve participants within each run for visualization. The heights of the bars represent the average consistency values across 20 runs in the discovery dataset, with error bars indicating the standard deviations.

**Figure 3.**
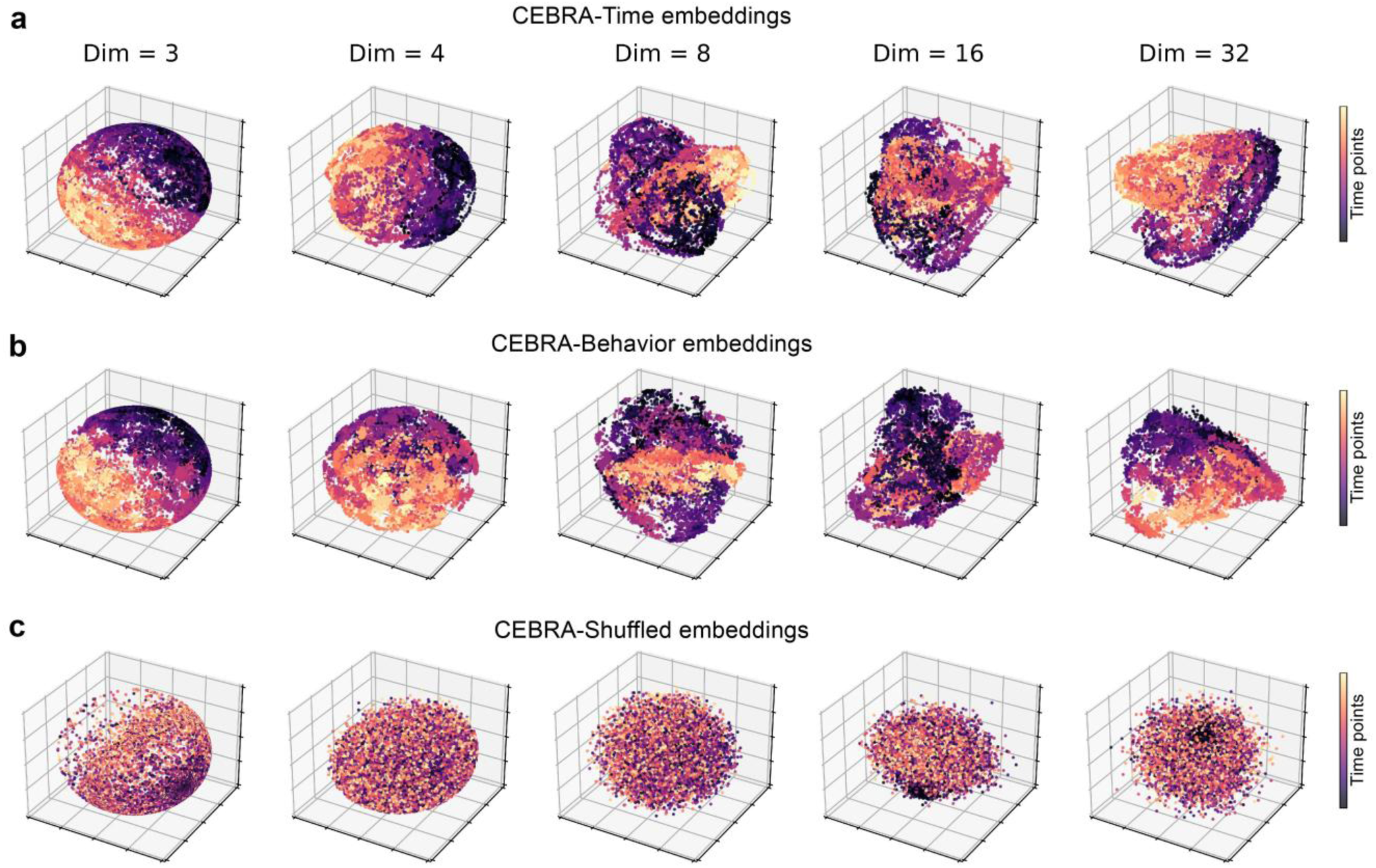
Neural trajectories in latent spaces derived from region-level MEG data of an example participant during story comprehension. **a-c)** The neural latent spaces were generated from a participant listening to the first story. Each latent space contained 10,145 CEBRA embeddings. We selected this example participant whose latent embeddings had the highest consistency with those of the other participants. Five different latent spaces with specific dimensions (three, four, eight, 16, and 32) were derived from CEBRA-Time, CEBRA-Behavior and CEBRA-Shuffled, respectively. For neural latent spaces with more than three dimensions, we selected the three latent components with the highest Shannon entropy for visualization.

Previous studies have shown that naturalistic stimuli can modulate neural activity with increased synchronization across individuals, presumably reflecting stimulus-driven responses^46–48^. If neural latent embeddings represent linguistic information from natural stories, they should be consistent across pairs of participants listening to the same story. To test this, we calculated the determination coefficient (*R*^2^) score of the linear regression between neural embeddings from pairs of participants (Fig. 2f). We then observed a significant interaction between technique and dimensionality (repeated-measures analysis of variance (ANOVA), *F*(12, 4750) = 537.391, *P* < 0.001, partial *η*^2^ = 0.576; Fig. 2g). We also found that CEBRA-Time significantly outperformed other benchmarks in producing consistent embeddings across participants in each dimension setting (post-hoc Tukey’s honest significant difference (HSD), *Ps* < 0.001). In addition, we found that the dimensionality of neural embeddings significantly influenced their consistency (post-hoc Tukey’s HSD, *Ps* < 0.01). Specifically, the linear model better fit the relationship between dimensionality and consistency in CEBRA-Time, as indicated by the lowest Akaike Information Criterion (AIC) and Bayesian Information Criterion (BIC) values (AIC = 472.401, BIC = 497.851; Supplementary Table 1). This indicated that the inclusion of more components led to a linear increase in the ability to represent shared information across participants during naturalistic story comprehension.

Furthermore, when neural embeddings corresponding to the same word are close together and form a stable representation of that word, this word representation should be decodable using these neural embeddings. To test it, we trained and evaluated the K-nearest neighbors (KNN) classifiers to decode unique words within each latent space, using neural embeddings based on the nested tenfold cross-validation scheme (Fig. 1c). We observed a significant interaction between technique and dimensionality on decoding accuracy (repeated-measures ANOVA, *F*(12, 4750) = 190.700, *P* < 0.001, partial *η*^2^ = 0.325; Fig. 4a). Neural embeddings derived from CEBRA-Time significantly outperformed those derived from the benchmarks in each dimension setting (post-hoc Tukey’s HSD, *Ps* < 0.001). In addition, dimensionality had a significant effect on the decoding accuracy with CEBRA-Time embeddings, as indicated by post-hoc tests (*Ps* < 0.001), except for the comparison between four and eight dimensions and the comparison between 16 and 32 dimensions (*Ps* > 0.05). However, dimensionality did not affect decoding accuracies with neural embeddings derived from the other techniques (post-hoc tests, *Ps* > 0.05). Moreover, we observed that the relationship between dimensionality and decoding accuracy with CEBRA-Time embeddings followed a logarithmic trend (Supplementary Table 1), where improvements in decoding accuracy became progressively smaller as the dimensionality increased.

**Figure 4.**
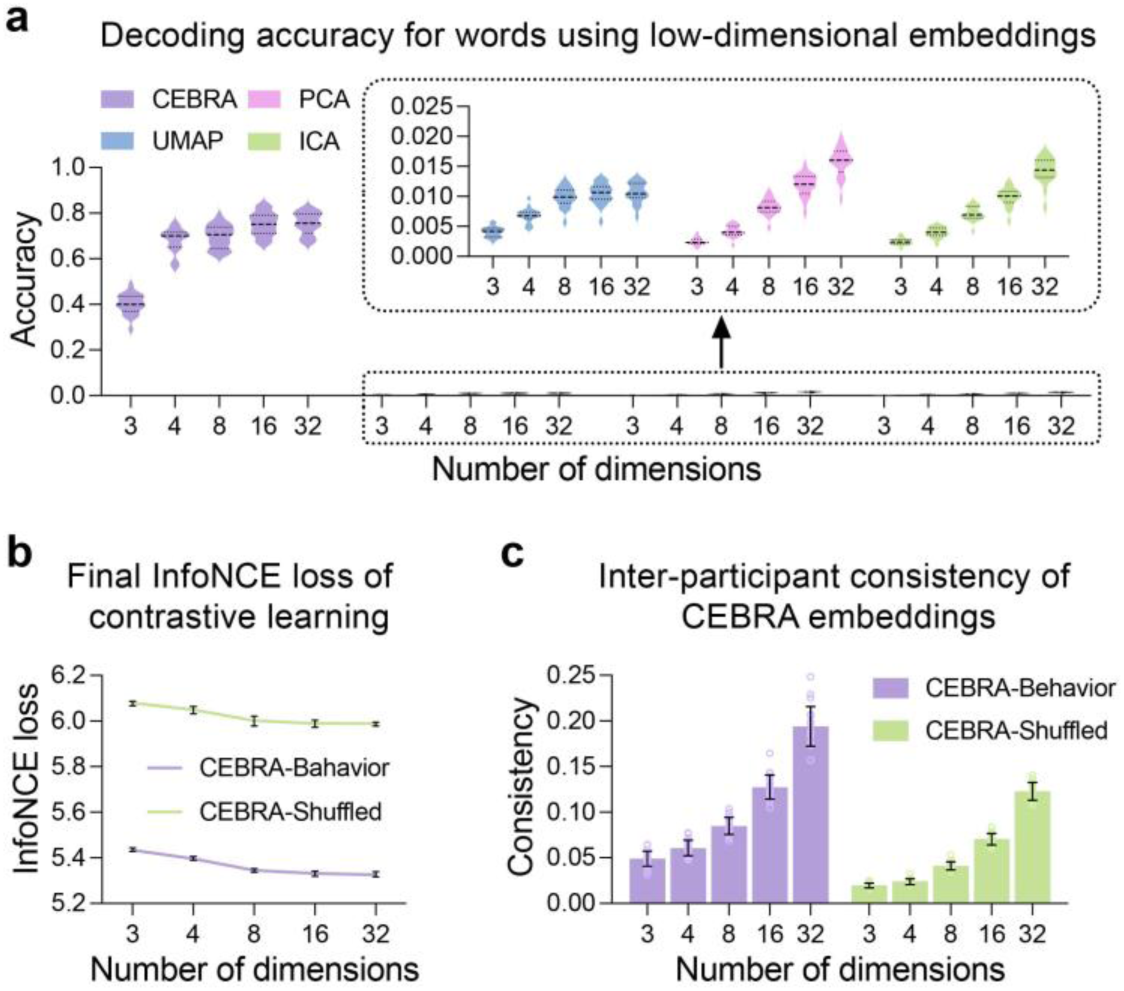
Exploration of language-relevant information in latent spaces. **a)** Word decoding accuracy using low-dimensional neural embeddings obtained from CEBRA-Time and other benchmark techniques. The adjusted balanced accuracies within each run in the discovery dataset (*n* = 20) were averaged to plot the violin plots. **b)** Final InfoNCE loss from the behavioral contrastive learning with lower values indicating better alignment between neural latent embeddings and contextual BERT embeddings. For visualization, averages were calculated across participants within each run. The purple and green lines indicate the average loss values across 20 runs, with black error bars indicating the range of loss values. **c)** The inter-participant consistency of the CEBRA-Behavior and CEBRA-Shuffled embeddings. The heights of the bars represent the average consistency values over these 20 runs, with the error bars indicating the standard deviations.

Taken together, we showed that the CEBRA-Time embeddings had higher consistency across participants and better discrimination of individual words compared to benchmarks, suggesting that the CEBRA framework excels at extracting meaningful components from high-dimensional neocortical responses during natural language comprehension. Furthermore, the shared information captured by CEBRA-Time directly benefits from higher-dimensional latent representations, whereas the linguistic information appears to be compressed into low-dimensional representations.

### Neural trajectories in latent spaces during natural language comprehension align with contextual activations of the pre-trained BERT model

Using self-supervised CEBRA-Time, we uncovered neural latent spaces from macroscale brain responses, providing a basic architecture for exploring neural representations of natural language. However, these latent spaces may contain both language-relevant and language-irrelevant information that contributed to consistent neural embeddings across participants. Here, we used behavioral contrastive learning with auxiliary labels to generate neural latent trajectories that were explicitly constrained by language-relevant features (Fig. 3b,c). These auxiliary labels were derived from activations of the last layer of a pre-trained version of bidirectional encoder representation from transformers (BERT) model, which captures contextual information from texts, including lexical content and syntactic relationships^19,49,50^. In addition, we used the loss function derived from behavioral contrastive learning to evaluate the alignment of neural latent embeddings with auxiliary labels represented by the contextual activations of the pre-trained BERT model.

We fit CEBRA-Behavior and its control models with a range of dimension settings, using the BERT embeddings or their temporally shuffled versions as auxiliary labels (see Methods). We then extracted the final Information Noise Contrastive Estimation (InfoNCE) loss from contrastive learning for both CEBRA-Behavior and -Shuffled^30^. We observed a significant interaction between auxiliary variable and dimensionality on InfoNCE loss (repeated-measures ANOVA, *F*(4, 2360) = 53.164, *P* < 0.001, partial *η*^2^ = 0.083; Fig. 4b). Specifically, the CEBRA-Behavior embeddings had significantly less final loss than CEBRA-Shuffled embeddings in every dimension setting (post-hoc Tukey’s HSD, *Ps* < 0.001). This indicated that CEBRA-Behavior embeddings better discriminated samples in auxiliary labels and showed better alignment with contextual BERT embeddings. Moreover, the final InfoNCE loss was significantly affected by the dimensionality in both types of latent embeddings (post-hoc tests, *Ps* < 0.001), except for the comparison between 16 and 32 dimensions in CEBRA-Shuffled (*P* > 0.05). We observed a logarithmic relationship between dimensionality and InfoNCE loss in CEBRA-Behavior (Supplementary Table 1), indicating that increasing the number of latent components improved the alignment between CEBRA-Behavior embeddings and contextual BERT embeddings, but the effect diminished with further increases.

We also examined the consistency of latent embeddings across participants. The interaction effect between auxiliary variable and dimensionality was significant (repeated-measures ANOVA, *F*(4, 2360) = 235.395, *P* < 0.001, partial *η*^2^ = 0.285; Fig. 4c). In addition, CEBRA-Behavior significantly outperformed CEBRA-Shuffled in producing consistent latent embeddings in each dimension setting (post-hoc Tukey’s HSD, *Ps* < 0.001), suggesting that CEBRA-Behavior embeddings better capture shared neural components across participants than those derived from control models. Furthermore, latent dimensionality significantly affected the consistency of CEBRA-Behavior embeddings (post-hoc tests, *Ps* < 0.001), with a linear relationship observed as the lowest AIC and BIC values in the first-order polynomial model (Supplementary Table 1).

In summary, the significant improvements in consistency and the reductions in final InfoNCE loss of CEBRA-Behavior latent embeddings, compared to their control counterparts, demonstrate their potential to capture shared, language-relevant neural components from large-scale neocortical responses during naturalistic story comprehension. Moreover, consistent with the results of CEBRA-Time, the amount of shared information captured by CEBRA-Behavior is associated with higher-dimensional latent representations, while the language-relevant information can be sufficiently represented using relatively low-dimensional latent spaces.

### Neural latent spaces contain a significant but subtle amount of information about part-of-speech tags and semantic categories

After using CEBRA-Time and CEBRA-Behavior models to unfold neural trajectories in latent spaces from human neocortical responses, we performed multivariate decoding to investigate how word-level latent embeddings in these latent spaces relate to candidate linguistic features. We first examined the amount of information contained in the latent spaces about two lexical features: part-of-speech tags and semantic categories. To do this, we trained and evaluated KNN classifiers to assess the extent to which these categorical features could be decoded using word-level latent embeddings derived from the CEBRA-Time and CEBRA-Behavior models (Fig. 5a,b). We found that the neural latent spaces of human neocortical responses contained a significant yet subtle amount of information about these categorical features.

**Figure 5.**
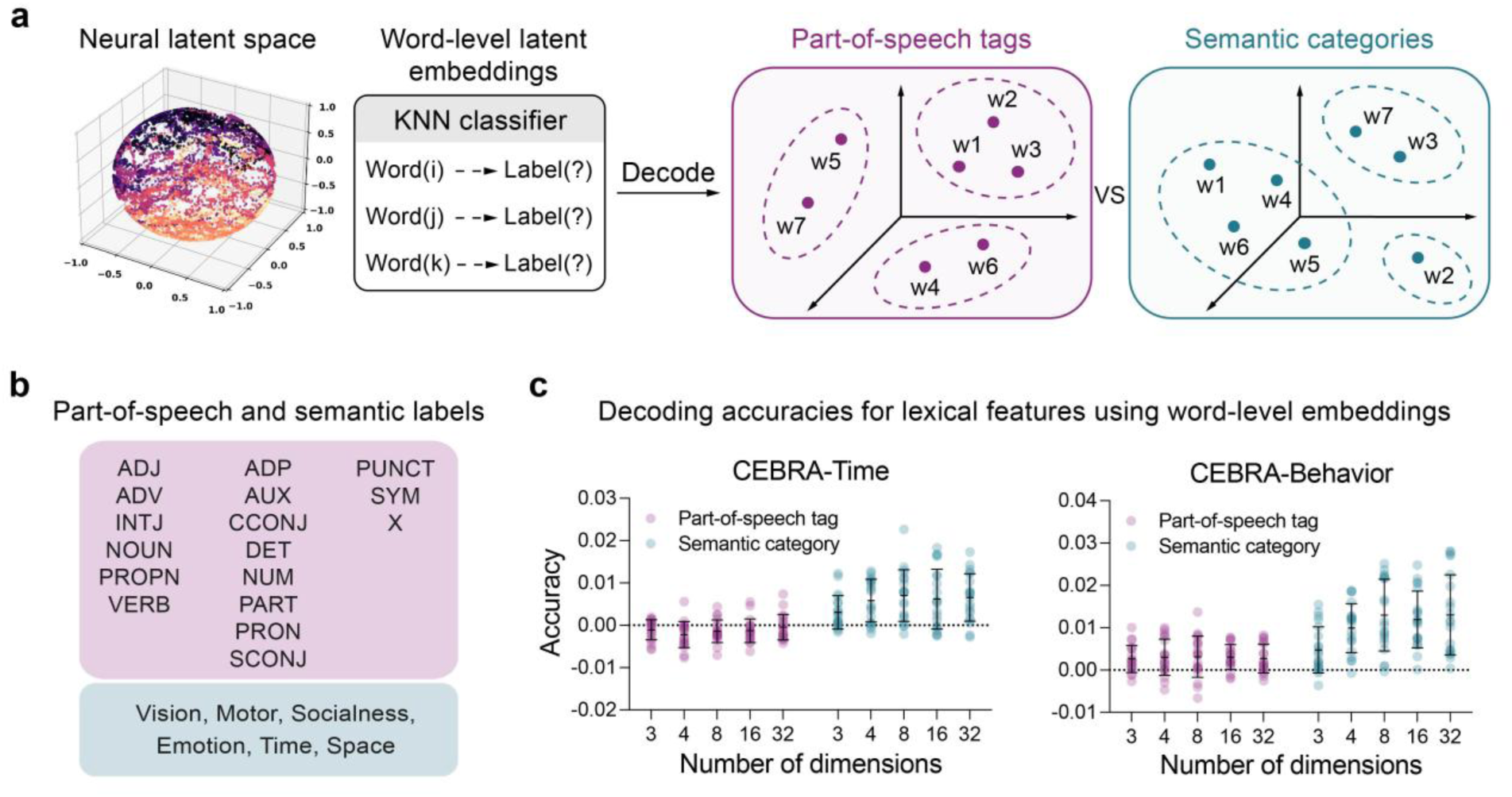
Multivariate decoding revealed the amount of information about part-of-speech and semantic categories in neural latent spaces. **a)** If word-level latent embeddings were organized according to part-of-speech tags or semantic categories in the latent space, word-level latent embeddings with the same tag or class should cluster together and can be decoded by a classifier. **b)** The universal part-of-speech tag system and six semantic categories were used as targets for KNN classifiers. **c)** Adjusted decoding accuracies for part-of-speech and semantic categories using word-level CEBRA embeddings from 20 runs in the analysis dataset. Each observation represents the average decoding accuracy within each run, calculated across test sets and then across participants. The black lines represent the means and the standard deviations.

For CEBRA-Time, we observed that part-of-speech decoding accuracies were significantly below chance levels (two-tailed randomization tests, *Ps* < 0.01), except for those using word-level latent embeddings of three dimensions (*P* = 0.055) and 32 dimensions (*P* = 0.360). In addition, decoding accuracies for semantic categories were significant in each dimension setting (*Ps* < 0.001; Supplementary Fig. 2). This showed that information about semantic categories was present in the latent spaces derived from CEBRA-Time models. We also observed a significant but minor interaction between attribute and dimensionality (repeated-measures ANOVA, *F*(4, 2360) = 4.076, *P* = 0.003, partial *η*^2^ = 0.007; Fig. 5c), and semantic decoding significantly outperformed part-of-speech decoding in each dimension setting (post-hoc tests, *Ps* < 0.001). We further observed that the mean-only model better fit the relationship between dimensionality and part-of-speech decoding accuracy (Supplementary Table 1), indicating a limited effect of dimensionality. Moreover, the logarithmic and mean-only models better fit the relationship between dimensionality and decoding accuracy for semantic categories with the lowest AIC and BIC values, respectively.

We further confirmed these results by investigating decoding accuracy with word-level latent embeddings derived from CEBRA-Behavior models. We observed that decoding accuracies for both part-of-speech tags and semantic categories were significant across all dimension settings (two-tailed randomization tests, *Ps* < 0.001; Supplementary Fig. 2). In addition, the interaction between attribute and dimensionality was significant but still small (repeated-measures ANOVA, *F*(4, 2360) = 10.805, *P* < 0.001, partial *η*^2^ = 0.018; Fig. 5c). We observed that semantic decoding was significantly better than part-of-speech decoding in each dimension setting (post-hoc tests, *Ps* < 0.05). Furthermore, we observed a logarithmic relationship between dimensionality and decoding accuracy for semantic categories with the lowest AIC and BIC values (Supplementary Table 1). This indicated that increases in dimensionality resulted in progressively smaller improvements in decoding accuracy for semantic categories. Again, the mean-only model better fit the relationship between dimensionality and decoding accuracy for part-of-speech.

Given the level of decoding accuracy for part-of-speech tags and semantic categories, these results suggest that neural latent spaces contain a statistically significant but subtle amount of information about the two lexical features. In addition, latent dimensionality appears to influence the extent to which semantic categories are captured, possibly in a logarithmic manner. However, the amount of information about part-of-speech tags may be irrelevant to latent dimensionality.

### Neural latent representations of individual words are primarily organized according to syntactic dependencies rather than lexical content

We then used representational similarity analysis (RSA) to investigate the correspondence of word-level embeddings in the latent spaces with distributed lexical representations or syntactic dependency relations. Briefly, we computed the representational distance matrix (RDM) using the cosine distance between word-level embeddings of CEBRA-Time and -Behavior, respectively. We then computed distributed lexical RDMs using static word embeddings and then created structured syntactic RDMs based on dependency relations of sentences (Fig. 6a). By evaluating the semi-partial correlations between neural latent RDMs and two linguistic RDMs, we found that unique linguistic contributions were significant in CEBRA-Time and -Behavior (two-tailed randomization tests, *Ps* < 0.001; Supplementary Fig. 3), suggesting that neural trajectories in latent spaces capture information about lexical representation and syntactic relations in naturalistic stories.

**Figure 6.**
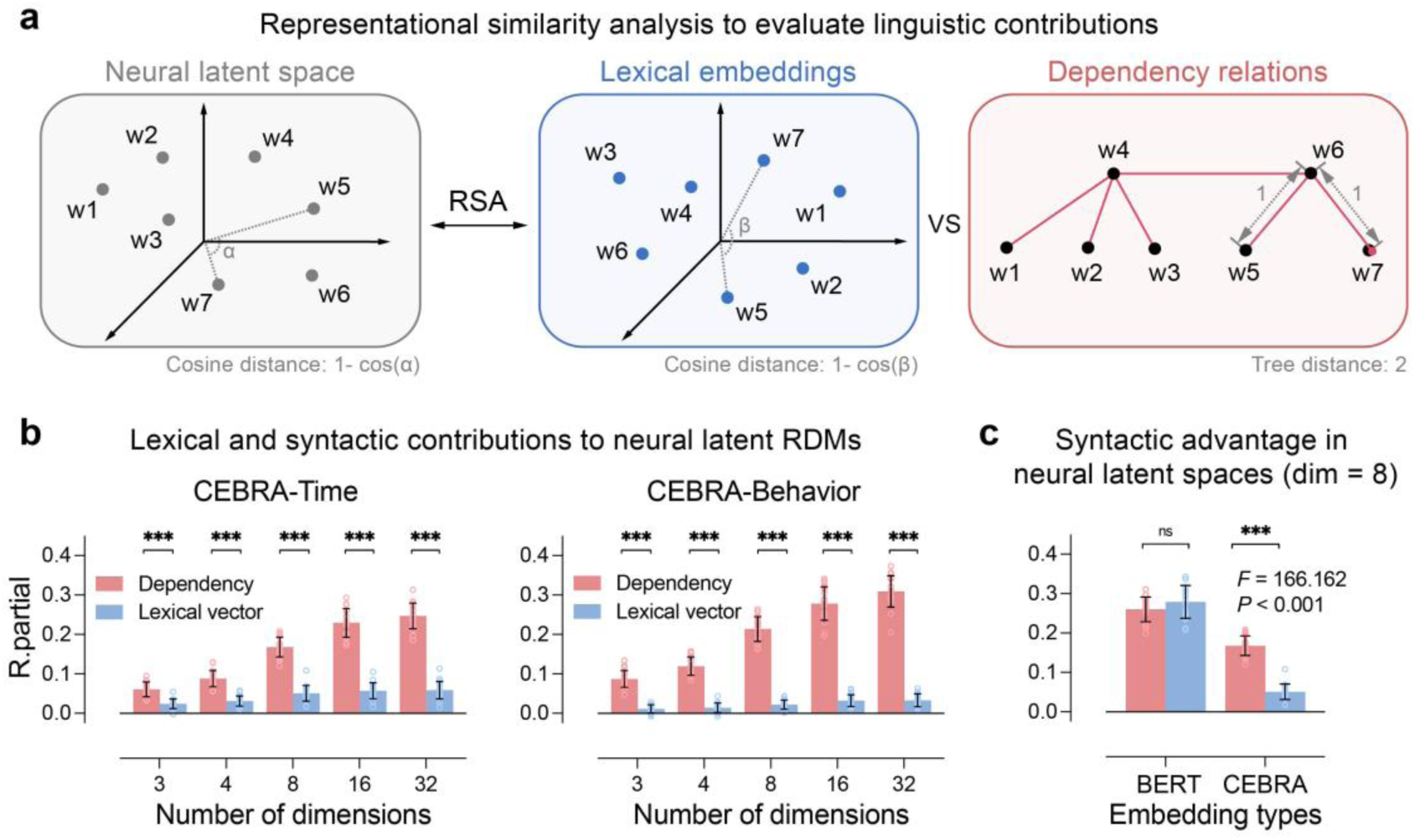
Representational similarity analysis revealed unique contributions of distributed lexical RDMs and structured syntactic RDMs to neural latent RDMs. **a)** Word-by-word neural, distributed, and structured RDMs of sentences can be computed from vector representations or syntactic dependency trees. The semi-partial Spearman correlation coefficients (partial *R*) were used to assess the unique contributions of distributed and structured RDMs to neural latent RDMs. **b)** Distributed and structured contributions to neural latent RDMs derived from CEBRA embeddings. For visualization, the partial *R* values were averaged across participants within each run in the analysis dataset (*n* = 20). **c)** Distributed and structured contributions to RDMs calculated from 8-D CEBRA-Behavior embeddings and contextual BERT embeddings, tested by the repeated-measures ANOVA. The heights of bars represent the average partial *R* values across 20 runs, while the error bars indicate the standard deviations. Triple asterisks (***) indicate p-values less than 0.001 and “ns” indicates p-values greater than 0.05 in the post-hoc tests.

In CEBRA-Time, we observed a significant interaction effect between linguistic feature and dimensionality (repeated-measures ANOVA, *F*(4, 2360) = 676.903, *P* < 0.001, partial *η*^2^ = 0.534; Fig. 6b). The structured syntactic RDMs explained significantly more variance in the neural latent RDMs than the distributed lexical RDMs in each dimension setting (post-hoc Tukey’s HSD, *Ps* < 0.001). We then observed logarithmic relationships between the dimensionality and the contributions of two linguistic features (Supplementary Table 1). Moreover, the contribution of structured syntactic RDMs was more sensitive to the dimensionality of latent spaces (linear mixed-effect model, standardized *β* = 0.968, SE = 0.015, *t*(1168) = 66.109, *P* < 0.001) than distributed lexical RDMs (*β =* 0.375, SE = 0.028, *t*(1168) = 13.444, *P* < 0.001). These results were subsequently confirmed in CEBRA-Behavior, where the interaction was also significant (*F*(4, 2360) = 548.532, *P* < 0.001, partial *η*^2^ = 0.482; Fig. 6b). Furthermore, the structured syntactic RDMs explained significantly more variance in the neural latent RDMs than the distributed lexical RDMs in each dimension setting (post-hoc tests, *Ps* < 0.001). We then found logarithmic relationships between the dimensionality and the contributions of linguistic features (Supplementary Table 1). Again, the contribution of structured syntactic RDMs was more affected by the latent dimensionality (*β* = 0.975, SE = 0.015, *t*(1168) = 66.350, *P* < 0.001) than distributed lexical RDMs (*β =* 0.498, SE = 0.025, *t*(1168) = 20.192, *P* < 0.001). The results of the multivariate analyses were validated using CEBRA-Time and -Behavior embeddings transformed from the nonlinear encoder with a different architecture (Supplementary Fig. 4-7 and Table 1).

Since CEBRA-Behavior embeddings were obtained through behavioral contrastive learning, one possibility was that the bias of the auxiliary variable led to the advantage of syntactic relations observed in CEBRA-Behavior embeddings. To test this, we computed word-by-word RDMs from the contextual BERT embeddings used in behavioral contrastive learning. Semi-partial Spearman correlations were computed to assess the unique contributions of distributed lexical RDMs and structured syntactic RDMs to the contextual BERT RDMs. We found that the contributions of syntactic relations differed between contextual BERT RDMs and 8-dimensional (8-D) CEBRA-Behavior RDMs, as indicated by a significant interaction (repeated-measures ANOVA, *F*(1, 19) = 166.162, *P* < 0.001, partial *η*^2^ = 0.897; Fig. 6c). The effect was replicated using neural latent embeddings across other dimension settings (Supplementary Fig. 1c). These results suggest that the advantage of syntactic relations in CEBRA-Behavior embeddings reflects the representational preference of the human neocortex, which cannot be directly explained by the bias of contextual BERT embeddings as auxiliary labels in behavioral contrastive learning.

Taken together, these results suggest that neural latent geometry is primarily associated with syntactic relations. Moreover, contrary to our initial hypothesis, the advantage of syntactic relations in latent space may not simply correspond to high-dimensional representations resulting from efficient coding, since this syntactic advantage shows up in neural latent embeddings of different dimensions. Rather, the representation of syntactic relations seems to be more compressive or robust compared to lexical content. Namely, neural trajectories in latent spaces may compress more information about syntactic relations than lexical features.

### The brain-inspired RC system with near-critical dynamics can better approximate observed neural trajectories in latent spaces

To investigate the mechanism underlying the observed neural trajectories that compact more information about syntactic relations compared to lexical features, we implemented brain-inspired reservoir computing (RC) systems^51,52^ to simulate human neocortical responses during natural language comprehension (Fig. 7a). Previous studies have shown that the human brain accumulates information over time and uses it to facilitate sequence processing, where the trace of past information can influence ongoing neural processes^53–55^. During language comprehension, the human brain also uses previous linguistic information to better predict and encode new incoming words^56,57^. These findings suggest that the memory capacity or integrative timescale of cortical responses may influence cortical dynamics when processing continuous linguistic stimuli. Given this, brain-inspired RC systems with good memory capacity may generate dynamics that more closely resemble observed neural trajectories with syntactic advantage. To test this, we configured the stability of the system’s dynamics to investigate how the global memory capacity contributes to the approximation of CEBRA-Time embeddings of 32 dimensions.

**Figure 7.**
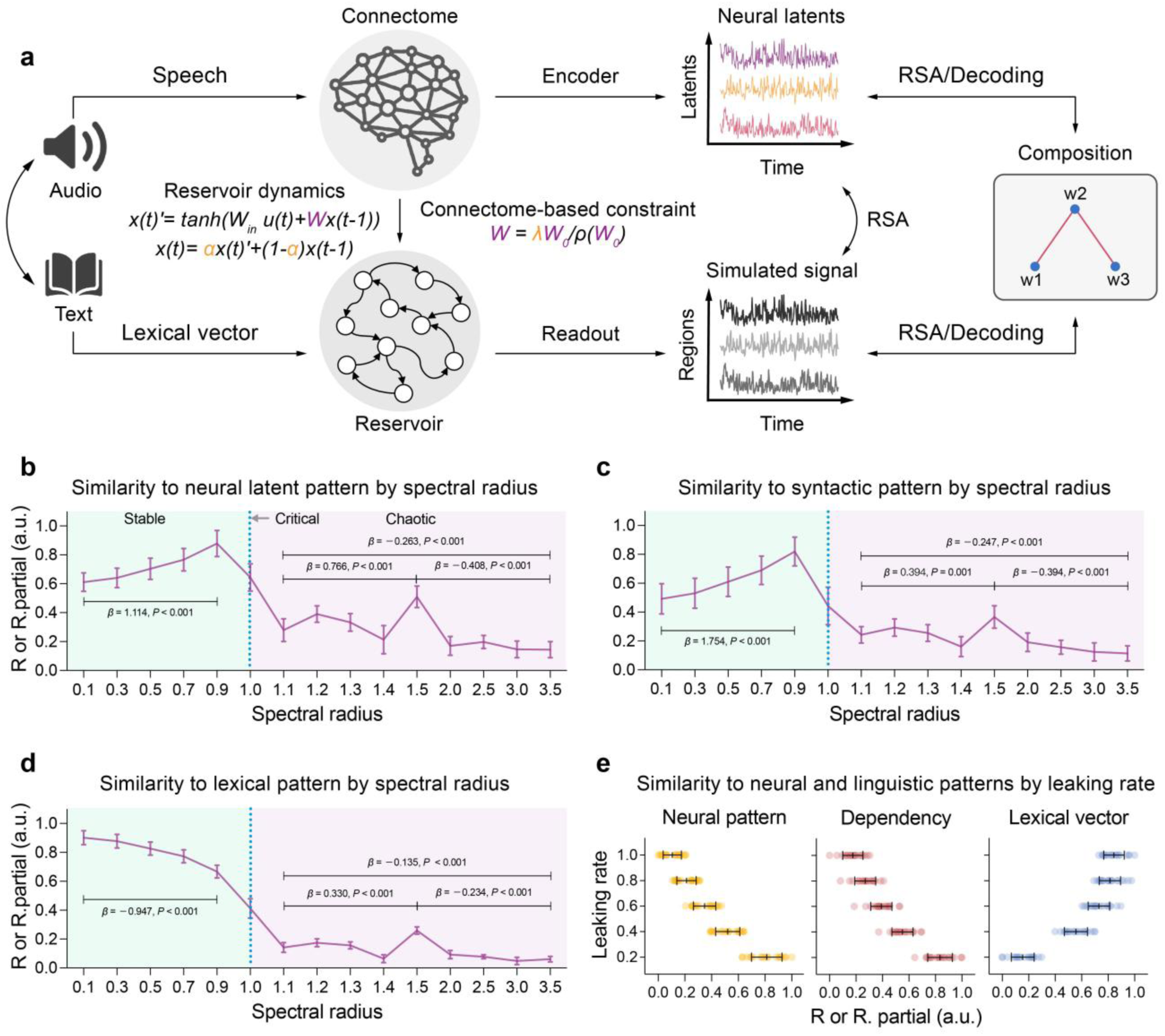
Simulation of observed neural trajectories using the brain-inspired RC system. **a)** The connectome-based reservoirs received temporally aligned lexical embeddings to generate activation states. We tuned the spectral radius (λ) to configure the memory capacity and adjusted the leaking rate (α) to configure the integration strategy of the brain-inspired RC system (see Methods; figure layout based on Lappalainen et al., 2024, under CC BY 4.0 license). **b-d)** The brain-reservoir similarity, the unique contribution of distributed RDMs, and the unique contribution of structured RDMs as functions of the spectral radius with the leaking rate fixed at 0.4, tested by linear mixed-effect models. The Fisher z-transformed (semi-partial) correlations were averaged within each run and then normalized over the spectral radii for visualization. The lines and bars represent the means and the standard deviations across 20 runs in the simulation dataset. **e)** The brain-reservoir similarity, the unique contributions of distributed and structured RDMs across leaking rates, with the spectral radius fixed at 0.9. Each observation represents the average score across participants within each run. The black lines represent the means and the standard deviations.

To configure the stability of dynamics, we adjusted the spectral radius, the largest absolute eigenvalue of the reservoir’s connection matrix. This parameter regulates the system’s ability to retain historical information by controlling the propagation of information along the connections. Theoretically, the system is in a stable state when the spectral radius is greater than zero and less than 1.0. Within this range, the larger the spectral radius, the slower the information decays, resulting in better storage capacity. When the spectral radius exceeds 1.0, information is continuously amplified along the connections, but the system enters a chaotic state. In this state, the system’s memory becomes unstable and unreliable, easily influenced by noise, the initial pattern, and previous inputs. At the critical point, where the spectral radius is about 1.0, the system reaches the “edge of chaos”, which is sensitive to external input and has a good memory capacity. After adjusting the spectral radius across the three dynamical regimes, we observed a higher brain-reservoir similarity in the stable state compared to the critical and chaotic states (Wilcoxon signed-rank test, *Ps* < 0.01; Supplementary Fig. 8g). Specifically, within the stable state, an increase in spectral radius approaching the critical point led to a steady improvement in brain-reservoir similarity (linear mixed-effect model, *β* = 1.114, SE = 0.049, *t*(1168) = 22.525, *P* < 0.001; Fig. 7b). These suggest that brain-inspired RC systems with near-critical dynamics, featuring sufficient memory capacity, can generate representations that more closely match observed neural trajectories.

We then performed multivariate analyses to investigate how the internal representations in brain-inspired RC systems relate to linguistic information. We observed that artificial internal RDMs were better related to distributed lexical RDMs and structured syntactic RDMs in the stable state compared to the other states (Wilcoxon signed-rank test, *Ps* < 0.001; Supplementary Fig. 8g). Similar effects were also observed in decoding accuracies for part-of-speech tags and semantic categories using internal activations of the brain-inspired RC systems (Supplementary Fig. 8b,e). In addition, within the stable state, we found distinct effects of spectral radius on different linguistic features. Specifically, the spectral radius positively affected the contribution of structured syntactic RDMs (linear mixed-effect model, *β* = 1.754, SE = 0.024, *t*(1168) = 73.884, *P* < 0.001, Fig. 7c). In contrast, we observed a negative effect on the contribution of distributed lexical RDMs (*β* = −0.947, SE = 0.013, *t*(1168) = −73.270, *P* < 0.001, Fig. 7d) and decoding accuracies for two lexical features (Supplementary Fig. 8a,d). These suggest that a superior memory capacity of the brain-inspired RC system specifically improves the representation of syntactic relations but impairs the representation of lexical features while the system is in the stable state.

In summary, we show that the brain-inspired RC systems in the stable state better approximate the observed neural trajectories while receiving continuous linguistic stimuli. Moreover, within the stable state, the brain-inspired RC systems closer to the critical point can produce more brain-like representations and adapt better to syntactic relations rather than lexical features.

### The brain-inspired RC systems, which favor the integration of past information, can produce more brain-like representations with syntactic advantages

Despite the historical information stored in the system, the immediate stimuli directly affect neural responses in the human brain. How to balance current and historical information in a system depends on the integration strategy used^58,59^. Compared to local lexical features, syntactic relations typically span multiple words and are more dependent on the sequence of input. Therefore, the human brain must effectively compress long-range and sequential information to construct compositional representations of natural language. Given this, we hypothesized that brain-inspired RC systems that prioritize the historical information in integration are more likely to generate internal representations that are related to observed neural trajectories. To test this, we adjusted the overall leaking rate of units in the reservoir to configure the system’s preference between current input and historical information. Theoretically, as the leaking rate decreases, the brain-inspired RC system reduces the weight of new inputs and increases its reliance on past states, thereby increasing its preference for historical information in the operation of integration.

We first evaluated the effect of the leaking rate on the similarity between neural latent RDMs and artificial internal RDMs from the brain-inspired systems with near-critical dynamics (spectral radius = 0.9). We found that the leaking rate negatively affected the brain-reservoir similarity (*β* = −0.802, SE = 0.015, *t*(1168) = −51.883, *P* < 0.001; Fig. 7e). This suggests that brain-inspired RC systems with a preference for historical information can generate more brain-like representations. In addition, we observed that the leaking rate also affected the contributions of linguistic RDMs in different manners. Specifically, increasing the leaking rate decreased the contribution of structured syntactic RDMs (*β* = −0.917, SE = 0.007, *t*(1168) = −123.564, *P* < 0.001) but increased the contribution of distributed lexical RDMs (*β* = 0.861, SE = 0.012, *t*(1168) = 71.587, *P* < 0.001). We also observed that the leaking rate had a positive effect on decoding accuracies for part-of-speech and semantic categories (Supplementary Fig. 8c,f). These findings suggest that the brain-inspired RC systems, with a preference for previous states, can produce internal representations that more closely match syntactic relations while showing weaker alignment with lexical features.

Furthermore, we examined the combined effect of spectral radius and leaking rate on the internal representations of brain-inspired RC systems. First, we stratified and averaged the data over the spectral radii into stable, critical, and chaotic states to focus on these dynamical regimes (see Methods). We then observed a significant interaction between leaking rate and stratified state on the brain-reservoir similarity (*F*(2, 275) = 248.578, *P* < 0.001, partial *η*^2^ = 0.644). This indicated that the effect of integration strategy on brain-reservoir similarity depended on the memory capacity.

In addition, we observed significant interaction effects on the contributions of structured syntactic RDMs (*F*(2, 275) = 198.026, *P* < 0.001, partial *η*^2^ = 0.590) and distributed lexical RDMs (*F*(2, 275) = 114.470, *P* < 0.001, partial *η*^2^ = 0.454, Supplementary Fig. 9). However, within the stable state, although the effects of spectral radius and leaking rate were significant for brain-reservoir similarity, lexical and syntactic contributions (repeated-measures ANOVA, *Ps* < 0.001, partial *η*^2^ ranging from 0.364 to 0.926), their interaction effects were not significant (*Ps* > 0.05), except for a subtle interaction effect on syntactic contributions (*F*(1, 477) = 5.080, *P* = 0.025, partial *η*^2^ = 0.011). These findings suggest that memory capacity and integration strategy significantly shape internal representations as the brain-inspired RC system transitions through the three dynamical regimes. While in the stable state, they independently influence the system’s internal representations.

Taken together, our simulation shows that the brain-inspired RC systems, with near-critical dynamics and favoring the integration of historical information rather than the immediate stimuli, can generate more brain-like representations specifically adapted to syntactic relations. This suggests that the storage and prioritization of long-range information may be two key computations for compacting information about syntactic relations into neural trajectories in latent spaces.

## Discussion

By examining neural trajectories in latent spaces, we investigated the how the human neocortical responses correspond to diverse linguistic information and flexibly support natural language composition. Using nonlinear dimensionality reduction, we generated neural trajectories in latent spaces from region-level MEG data of participants listening to 60 naturalistic stories with a total duration of approximately six hours. We demonstrate that these trajectories are language-relevant by showing that they capture consistent components across participants, discriminate single-word representations, and align with activations of the pre-trained bidirectional encoder representation from transformers (BERT) model. We then reveal the associations of lexical and syntactic features with neural latent geometry using multivariate decoding and representational similarity analysis (RSA). We further show that the human cortex orchestrates distinct coding strategies to facilitate the composition of natural language, in which syntactic relations are associated with cortical coding with more efficient compression compared to lexical features. Furthermore, our simulation using brain-inspired RC systems demonstrates that observed neural trajectories can be simulated with near-critical dynamics and a preference for historical information. Taken together, these findings offer a new perspective on the neurocomputational mechanisms underlying natural language composition and challenge views that place primacy on lexical content. They also provide insights for the development of brain-inspired intelligent systems capable of more effectively integrating long-range and contextual information in the real-world language process.

In this study, we used a dimensionality reduction technique called CEBRA^30^, which can generate neural trajectories in latent spaces from multivariate neural data. This nonlinear technique allows more effective identification of underlying structure from high-dimensional neural recordings than traditional linear techniques^30,60,61^ and allows us to determine how human language is related to large-scale and coordinated neocortical responses. Previous studies have revealed language-related responses in the human brain, with widespread areas showing neural synchronization between individuals during the processing of real-world stimuli such as movies or stories^46–48^. Associations between stimulus-driven cortical responses and activations from large language models (LLMs) have also been demonstrated^62–65^. We go beyond these approaches by quantifying the intersubject consistency of neural trajectories and examining the alignment of these trajectories with contextual BERT activations. Using high temporal resolution MEG data, we were able to precisely track the location of neural states in latent spaces, allowing multivariate analyses to uncover the relationship between neural states and linguistic features of naturalistic stories.

Our finding of a language-relevant latent space contributes to a growing body of literature identifying latent structures from the coordinated activation of neural populations in non-human animals. Electrode recording or two-photon calcium imaging has revealed low-dimensional latent spaces from neural activity in the hippocampus^28,66^, prefrontal cortex^67,68^, visual cortex^69,70^, and thalamic nucleus^71,72^. Similar low-dimensional architectures have also been identified in functional neuroimaging of the human brain under resting and task conditions^73–76^. Recent studies have examined the relationships between naturalistic stimuli and large-scale brain dynamics^77–80^; however, they focused on specific types of features and did not examine the construction of complex meanings. Using multivariate decoding and RSA, our study went beyond previous work by uncovering the integration of lexical content and syntactic relations in latent spaces. Our findings show that humans use similar low-dimensional architectures underlying the composition of natural language as those observed in non-human animals when engaged in other cognitive tasks.

We evaluated the different linguistic features in natural language composition and found that neural latent trajectories primarily reflect the composition based on syntactic relations. Previous imaging studies of language have suggested that lexical features may be sufficient to identify neural substrates for the language process, potentially indicating a central role of lexical content^5^. By examining neural latent patterns, we challenge these findings by showing that the representation of lexical content is less important than syntactic relations in neural latent spaces. Our work is consistent with recent research that has emphasized the structure-based process during language production and comprehension^81–84^. It is also supported by the research showing that semantically implausible but syntactically well-formed stimuli can fully engage the language network to an extent similar to intact sentences^85^. In addition, we observed that part-of-speech tags and semantic categories had weaker representations in latent spaces, indicating that they may not be central to neocortical responses during naturalistic story comprehension^85–87^. These findings suggest that neural trajectories in latent spaces arise primarily from the structure-based construction of meaning, rather than from a word-by-word traversal process based on lexical content. Future studies could manipulate semantic ambiguity or syntactic complexity to investigate how neural trajectories are causally altered by different linguistic conditions. It would also be valuable to compare neural trajectories across languages to investigate their resilience to cross-linguistic diversity^88,89^.

Research in human and non-human primates has shown that dimensionality varies across the human cortex^90,91^, adapts flexibly to task demands^16,42,92^, and changes during learning^43,70,93^. The current study shows that the relationship between natural language and the dimensionality of latent spaces depends on linguistic features, providing new insights into the cortical coding of language cognition^94,95^. We found a linear relationship between the latent dimensionality and the shared information encoded by individuals listening to the same story. However, the relationship between dimensionality and neural representations became logarithmic when we focused on language-related information. This logarithmic relationship suggests that an optimal number of latent components can robustly capture linguistic features. In particular, the syntactic relations are more sensitive to dimensionality than other features, as evidenced by higher growth rates in logarithmic models. These results, together with evidence highlighting the larger information of syntactic relations in latent spaces, demonstrate that neural latent spaces compress syntactic relations more efficiently than lexical content. This efficient compression, which differs from the typical coding strategies we hypothesized, implies compact neural components associated with information about syntactic relations and indicates a high efficiency of syntactic representations in the human neocortical activation^92,93^. Taken together, these findings suggest that the human neocortex coordinates multiple coding strategies to flexibly handle different linguistic features under resource constraints, which may distinguish human communication from that of other species^96^.

To further understand the computational mechanisms underlying observed neural trajectories during natural language processing, we used connectome-based reservoir computing (RC), a type of artificial neural network with biological constraints, to simulate human neocortical dynamics. This approach allowed us to adjust parameters and test their effects on the systems’ representations^97^. Our simulation shows that memory capacity and integration strategy jointly influence internal representations. Previous studies have shown that the human brain continuously accumulates input signals and uses historical information to facilitate the prediction and encoding of upcoming stimuli^53–55^, suggesting that previous information stored in the brain may influence the neural state at a given time. In this study, we investigated the effect of memory capacity on observed neural trajectories. We demonstrate that the brain-inspired RC system is better able to approximate observed neural trajectories and adapt to syntactic relations by retaining more historical information when in the stable state and approaching critical dynamics. In addition to the historical information, the current input can directly modulate the system’s states^58,59^. This balance between current input and past information in a system depends on the integration strategy. This applies to language, as recent research has shown that syntactic recognition emerges from neural circuits with a recurrent pathway that can integrate current input and past information^98,99^. We extend these findings by showing that brain-inspired RC systems can better simulate observed neural trajectories and specifically adapt to syntactic relations when they give more weight to past states in information integration. More generally, our work suggests that an intelligent system must contextualize information over time to effectively adapt to changing environments.

Future studies could combine other neuroimaging techniques with enhanced spatial resolution, such as intracranial and single-neuronal recordings^100^, to further validate the geometric organization of population activity in language-selective areas. This would provide computational insights into language cognition at the microscale level, which is not effectively assessed by non-invasive neuroimaging modalities. In addition, exploring neural latent structures in other high-order cognitive domains can determine whether there are similar or specific latent structures across different contexts. For example, recent research has shown that the human brain uses basis functions to compress social information^101^, indicating a universal scheme of neural coding across cognitive domains. Other studies have shown that neural trajectories differ across task conditions, showing the specificity of neural latent geometry^73,74^. Our findings on how the human neocortex performs language composition via neural latent trajectories may also have implications for clinical practice. Disruption of neural trajectories in latent spaces may be a potential biomarker of impaired language function^102,103^. If so, atypical neural trajectories in latent spaces would be detected in individuals with clinical conditions while they engage in naturalistic story comprehension.

In conclusion, we have investigated how the human brain flexibly coordinates cortical coding to composite natural language via neural trajectories in the latent spaces. We show that syntactic relations play a dominant role through more efficient compression than lexical content. This representational advantage of syntactic relations in neural latent spaces specifically benefits from the storage and prioritization of long-range information. Our work challenges current arguments that syntactic relations are not uniquely encoded by the human brain, distinct from lexical content.

## Methods

### SMN4Lang

The data used to prepare this work were obtained from the synchronized multimodal neuroimaging dataset for the study of brain language processing (SMN4Lang)^44^. This is a publicly available dataset containing the magnetoencephalography (MEG), diffusion magnetic resonance imaging (MRI), T1-weighted and T2-weighted MRI data from twelve participants (four females and eight males). During MEG acquisition, these participants listened to 60 naturalistic stories in Mandarin (ranging from four to seven minutes per story), for a total of nearly six hours of unique stimuli. We randomly divided the MEG recordings of 60 runs (stories) into three equal parts: one part for evaluating different dimensionality reduction methods (discovery dataset), another for analyzing neural trajectories in latent spaces using multivariate approaches (analysis dataset), and the third for computational modeling to investigate computational mechanisms (simulation dataset). Thus, each data subset consisted of 20 runs, representing nearly two hours of naturalistic stimuli.

### Data preprocessing

**MRI.** Raw MRI data were downloaded from the SMN4Lang. Minimal preprocessing pipelines from the Human Connectome Project (HCP) were then used to preprocess the structural and diffusion images^104^. Briefly, the structural preprocessing steps included gradient distortion correction, readout distortion removal, bias field correction, registration to the MNI standard space, down-sampling, and surface file registration. Diffusion preprocessing was then performed by normalizing the b0 image intensity across runs, correcting for susceptibility, eddy-current distortions, and motion, registering the diffusion data to the structure, and masking the data with the final brain mask.

**MEG.** The SMN4Lang provided the preprocessed MEG data at the sensor level. The preprocessing was performed in the Maxfilter (Elekta-Neuromag) and the MNE-Python^105^. The steps included magnetic artifact suppression using the temporal signal space separation, automatic identification and exclusion of bad channels using a default threshold value and interpolation method, identification and removal of ocular artifacts related to blinks, heartbeats, and eye movements using ICA, and finally application of a bandpass filter (0.1 to 40 Hz) to the denoised data.

### Annotations of transcripts

**Speech-to-text alignment.** Word boundaries in the audio were identified and aligned with the transcripts using the Montreal Forced Aligner (MFA) software (https://montreal-forced-aligner.readthedocs.io/), which uses the Kaldi ASR toolkit to perform forced alignment. In addition, before alignment, we corrected words in the transcripts, such as years (e.g., in 2021) and mathematical symbols (e.g., percentage), based on their actual pronunciation in Mandarin. After alignment, the MFA software provided the onset and offset time of each word in the audio.

**Static and contextual word embeddings.** Contextual 768-D word embeddings were obtained from a pre-trained BERT model (https://huggingface.co/google-bert/bert-base-chinese). This model contains 12 layers of bidirectional-attention transformer modules stacked on top of a non-contextual embedding layer. The transcript was formatted, divided into sentences, and tokenized using the model’s default tokenizer. Each sentence was then run through this pre-trained BERT model, and the activation of each word in the sentence was extracted. Static 100-D word embeddings were obtained using a pre-trained Word2Vec model (https://ai.tencent.com/ailab/nlp/en/embedding.html). **Part-of-speech tags, dependency relations, and semantic categories.** Part-of-speech tags and syntactic dependency relations were extracted using Stanza (https://stanfordnlp.github.io/stanza/). For the semantic categories, we trained a ridge classifier with static Word2Vec embeddings to predict six major semantic categories from 17,940 commonly used Chinese words in a large dataset of semantic ratings^106^. The semantic categories included vision, motor, socialness, emotion, time, and space, which were determined by the rating scores in this large dataset. The regularization parameter of the ridge classifier was searched among ten values log-spaced between 10^−1^ and 10^8^, based on a tenfold cross-validation framework. Finally, the trained classifier was used to predict the semantic categories for each word in naturalistic stories from SMN4Lang.

### Neural latent space modeling

**Source time courses.** Source estimation from sensor-level MEG data was performed in MNE-Python. The source space was modeled by a cortical mesh consisting of 8,196 vertices. Sensor positions were registered to T1-weighted structural images for each participant by aligning fiducials and the digitized head shape to the outer scalp. Forward models were then generated based on single-shell models. The epoch was extracted with the auditory delivery delay corrected (39.5 ms), which was then aligned with the audio’s onset and extended to the offset. The baseline correction was applied to the epoch data by subtracting the mean activity averaged from −2.0 s to 0 s relative to the audio’s onset. The source time courses of 8,196 vertices were constructed using the least-squares minimum norm method and then parcellated to the Schaefer-400 atlas^107–110^.

**Self-supervised CEBRA model.** Self-supervised CEBRA-Time models were used to generate neural trajectories in latent spaces from the parcellated MEG data^30^. First, model architectures and time offsets for contrastive learning were selected based on a grid search (Supplementary Fig. 1a), and the optimal architecture with the specific time offset showing the least InfoNCE loss was selected for subsequent analyses. We then applied a nonlinear transformation to each participant’s MEG data for each run through a feature encoder (selected model: offset40-model-4x-subsample), and performed time contrastive learning (selected time offset = 10) to generate latent embeddings by attracting similar (positive) samples and repelling dissimilar (negative) samples, using InfoNCE as the loss metric^102^. We also set a sufficient number of iterations to ensure that the loss converges. In addition, we explored three, four, eight, 16, and 32 output dimensions and kept other parameters unchanged to explore the effect of the dimensionality of latent spaces. Finally, three dimensionality reduction techniques, including PCA, ICA, and UMAP, were also applied to transform the parcellated MEG data into low-dimensional neural embeddings of three, four, eight, 16, and 32 dimensions. We also reported results using feature encoders with a different architecture (model: offset10-model, selected time offset = 10) in the Supplementary Material.

**Supervised CEBRA model.** The CEBRA-Behavior model was used with contextual BERT embeddings as auxiliary labels. First, the activation of the last layer of the BERT model was extracted. PCA was then applied to the contextual BERT embeddings for each run, with 100 components explaining over 70% of the variance in the original data and ensuring feature independence. The 100-D embeddings were then up-sampled to match the sampling rate of the source-level MEG data and to match the onset and duration of each word. A grid search was then performed to select the optimal model architecture and the time offset (Supplementary Fig. 1b). Next, feature encoders (selected model: offset40-model-4x-subsample) were used to apply a nonlinear transformation to the parcellated MEG data. Behavioral contrastive learning (selected time offset = 10) was performed to generate neural latent embeddings using 100-D BERT embeddings as auxiliary labels. In addition, three, four, eight, 16, and 32 output dimensions were also investigated. Finally, we used CEBRA-Shuffled models as controls, where the auxiliary labels were time-shuffled to ensure that they were not correlated with the neural signal.

**Consistency and InfoNCE loss.** To measure the consistency of neural latent embeddings across participants under the same naturalistic stimuli, we calculated the *R*^2^ value of the linear regression between latent embeddings from each pair of participants^30^. The *R*^2^ values were then averaged across pairs within each participant for statistical tests. In addition, the final values of the InfoNCE loss values were extracted from behavioral contrastive learning and used to measure how well neural latent embeddings discriminated positive, negative, and reference samples in auxiliary labels, reflecting their alignment with contextual BERT embeddings.

### Multivariate decoding analysis

**Decoding words using latent embeddings.** We used the KNN classifier with cosine distance metric to decode individual words using neural embeddings^112^. Based on time stamps, word labels were assigned to neural embeddings derived from CEBRA-Time, UMAP, PCA and ICA. Then, the nested tenfold cross-validation scheme was used to find the most optimal number of neighbors over the range [5, 10, 15, 20, 25] in the inner loop and to evaluate the decoding performance with the balanced accuracy score in the outer loop. The balanced accuracy score was adjusted so that 0 represented the random performance and 1 represented the perfect performance. Finally, the adjusted decoding accuracies were averaged across test sets within each participant and each run to obtain the final classifier performance.

**Decoding part-of-speech tags and semantic categories.** First, we averaged the neural latent embeddings derived from CEBRA-Time and CEBRA-Behavior within the period of each word to obtain word-level latent embeddings. Then, we used the KNN classifier with cosine distance metric to decode lexical part-of-speech tags and semantic categories using the word-level latent embeddings in each latent space. As in the case of word decoding, a nested tenfold cross-validation scheme was used to find the most optimal number of neighbors over the range [5, 10, 15, 20, 25] in the inner loop and to evaluate the performance in the outer loop. Finally, the adjusted decoding accuracies were averaged across test sets within each latent space to obtain the final performance for two lexical features, respectively.

### Representational similarity analysis

**Neural latent RDMs.** Each entry in the neural latent RDM was computed based on the cosine distance between the word-level latent embeddings. A between-participants approach was used to compute the cosine distances of one participant and the group-averaged word-level embeddings of the other participants from the same run^113^. For each participant and each sentence, the word (of one participant) by word (group-averaged) cosine distance matrix was first obtained. These matrices were then averaged with their transposes to ensure that the neural latent RDMs were symmetrical. The upper triangular parts of these RDMs were used to compute correlations with the upper triangular parts of the linguistic RDMs from the corresponding sentences (see details below).

**Distributed lexical RDMs and structured syntactic RDMs.** Two linguistic RDMs were computed between all word pairs for each sentence. The static Word2Vec embeddings and one-hot vectors representing part-of-speech tags were combined, resulting in 117-D lexical embeddings. The distance between each pair of words in the distributed lexical RDMs was measured using the cosine distance of the lexical embeddings. In addition, the structured syntactic RDM was derived from the dependency relationship of each sentence. Specifically, the distance between each pair of words was measured as the number of edges between them in the syntactic tree structure.

In addition, acoustic feature, word length, and word frequency were used as covariates. Acoustic RDMs were computed using the cosine distance of embeddings derived from a pre-trained Wav2Vec2 model (https://huggingface.co/facebook/wav2vec2-large-xlsr-53)^114^. The rate and length of each word were also used to compute difference values in their RDMs. Next, the upper triangular parts of the linguistic RDMs were extracted and then used to compute semi-partial Spearman correlation coefficients with the upper triangular parts of the neural latent RDMs, after removing the effects of acoustic feature, word length, word frequency, and the other linguistic feature. Finally, these semi-partial correlations were Fisher z-transformed and averaged within each run for statistical tests^115^ excluding sentences in the top and bottom 5% of length.

### Brain-inspired RC system

**Structural network construction.** The bidirectional nonlinear transformation between structural space and standard space was applied using the FMRIB Software Library (FSL)^116^. Then, 400 cortical regions^108^ and 16 subcortical regions^117,118^ in atlases were selected as seed regions to guide probabilistic tractography tracking as implemented in the ProbtrackX GPU program^119^. The network mode was used to compute a region-by-region matrix to quantify the number of streamlines seeded in one region that reached another. In addition, the average length of the streamline between the two regions was calculated and other parameters were configured according to default settings. Finally, the region-by-region matrix was normalized by the mean streamline length and mean voxel number of two regions to obtain the structural network for each partcipant^120^.

**Connectome-based reservoir computing.** We used connectome-based reservoir computing to emulate human neocortical responses during natural language comprehension, implemented with the BrainPy^121^. The reservoir computing system generally includes an input, a reservoir, and a readout layer. In this study, an input layer received and transmitted input signals (117-D lexical embeddings) to the reservoir. To simulate background noise, random Gaussian noise was added to the input signals. Then, the reservoir (i.e., the structural brain network) received these signals in a distributed manner and performed a nonlinear projection to emulate neocortical responses under naturalistic stories. Finally, a readout layer extracted the activations of target nodes in the reservoir (400 cortical regions) for multivariate analysis. The dynamics of the connectome-based reservoir was represented by serial vectors of activation states governed by the following equations^122^:

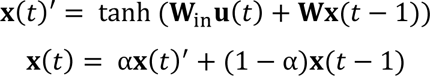

The **W**_in_ was the input matrix. The **u**(*t*) represented the input signal at time *t*. The **x**(*t*) was the activation state of all nonlinear units at time *t*. The α was the leaking rate. The **W** denoted the connection weights of the reservoir, defined by the structural brain network of each participant.

We investigated how memory capacity and integration strategy affect the internal representations of the brain-inspired RC system. We configured the memory capacity by changing the stability of the reservoir dynamics. Specifically, we scaled the spectral radius, defined as the maximum absolute eigenvalue of the **W**, by multiplying a tuning factor (λ) with 15 possible values [0.3, 0.5, 0.7, 0.8, 0.9, 1.0, 1.1, 1.2, 1.3, 1.4, 1.5, 2.0, 2.5, 3.0, 3.5]^51^. The RSA and decoding measures were stratified into stable (spectral radius = [0.3, 0.5, 0.7, 0.8, 0.9]) and critical (spectral radius ≈ 1.0) states. To distinguish between the critical and chaotic states, we characterized the chaotic state by selecting spectral radii between 1.5 and 3.5 (spectral radius = [1.5, 2.0, 2.5, 3.0, 3.5]). The results for spectral radii between 1.1 and 1.5 are also available in the Supplementary Material. Finally, to adjust the integration strategy, we tuned the leaking rate with five possible values [0.2, 0.4, 0.6, 0.8, 1.0].

### Statistics

Linear mixed-effect models were implemented to examine the effects of technique, auxiliary variable, linguistic feature, and dimensionality, with participant and run treated as random effects. Akaike Information Criterion (AIC) and Bayesian Information Criterion (BIC) were used to evaluate three candidate models characterizing the relationship between dimensionality and information representation in the latent spaces. In addition, randomization tests were performed to determine whether adjusted decoding accuracies and semi-partial correlation coefficients were significantly higher than chance. The null distributions were generated by randomly shuffling the target labels or neural latent RDMs 1,000 times. Moreover, to reduce the statistical complexity, we averaged data across participants within each run while examining the combined effect of spectral radius and leakage rate on the internal representations of brain-inspired RC systems. Wilcoxon rank-sum tests were used to compare the RSA and decoding measures between the three stratified states.

## Data availability

The synchronized multimodal neuroimaging dataset for studying brain language processing is publicly available on OpenNeuro (<u>10.18112/openneuro.ds004078</u>). The six semantic dimension database is available on OSF (<u>10.17605/OSF.IO/N5VKE</u>). Icons used in the figures were created by the authors <u>Freepik</u>, <u>Pixel perfect</u>, <u>Bharat Icons</u>, <u>Imaginationlol</u>, <u>Us and Up</u> from www.flaticon.com.

## Code availability

Structural and diffusion images were preprocessed using the minimal preprocessing pipelines of HCP^104^. Structural brain networks were constructed using FSL^116^ and the ProbtrackX GPU program^123^. The MEG data were analyzed using MNE-Python^105^. Neural trajectories in latent spaces were extracted using CEBRA^30^. Reservoir computing was performed with BrainPy^121^. Multivariate analyses were performed with scikit-learn^112^ and Pingouin^115^. Statistical tests were implemented using the lme4^124^, lmerTest^125^ and effectsize^126^ packages in R (version 4.4.1)^127^.

## Acknowledgements

This work was supported by the National Natural Science Foundation of China (NSFC: 32471100, 32371141). The authors thank Jiaqi Ma and Ziyi Jin for checking the annotation of transcripts.

## Supplementary Materials

**Supplementary Figure 1.**
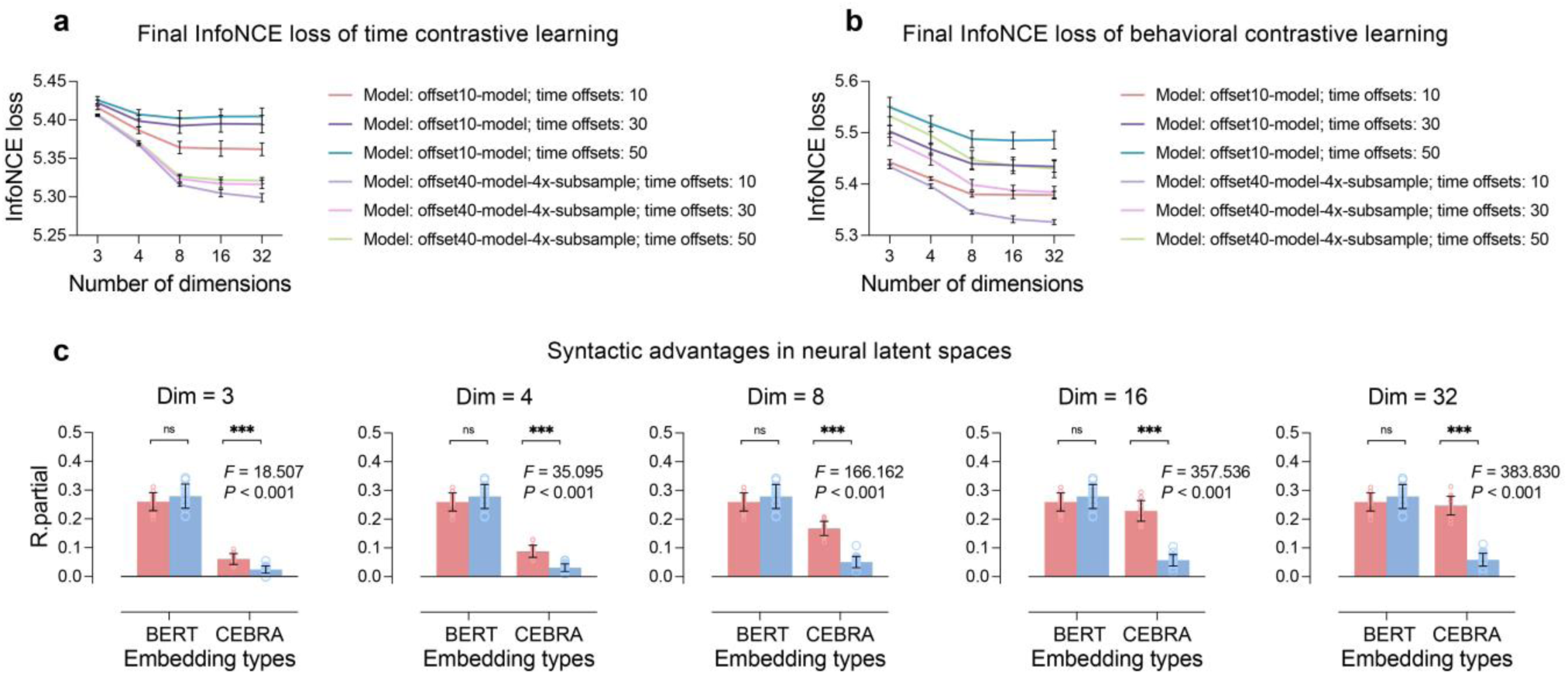
Grid search of hyperparameters in CEBRA models and syntactic advantages in contextual BERT RDMs and neural latent RDMs. **a-b)** Grid search was performed to find optimal model architectures and time offsets for contrastive learning with the lowest InfoNCE loss. The colored lines indicate the average loss values across 20 runs in the discovery dataset, with black error bars indicating the standard deviations. c) Distributed and structured contributions to RDMs computed from CEBRA-Behavior and contextual BERT embeddings, tested by repeated-measures ANOVA. The heights represent the average partial *R* values across 20 runs in the analysis dataset, while the error bars indicate the standard deviations. Triple asterisks (***) indicate p-values less than 0.001 and “ns” indicates p-values greater than 0.05.

**Supplementary Figure 2.**
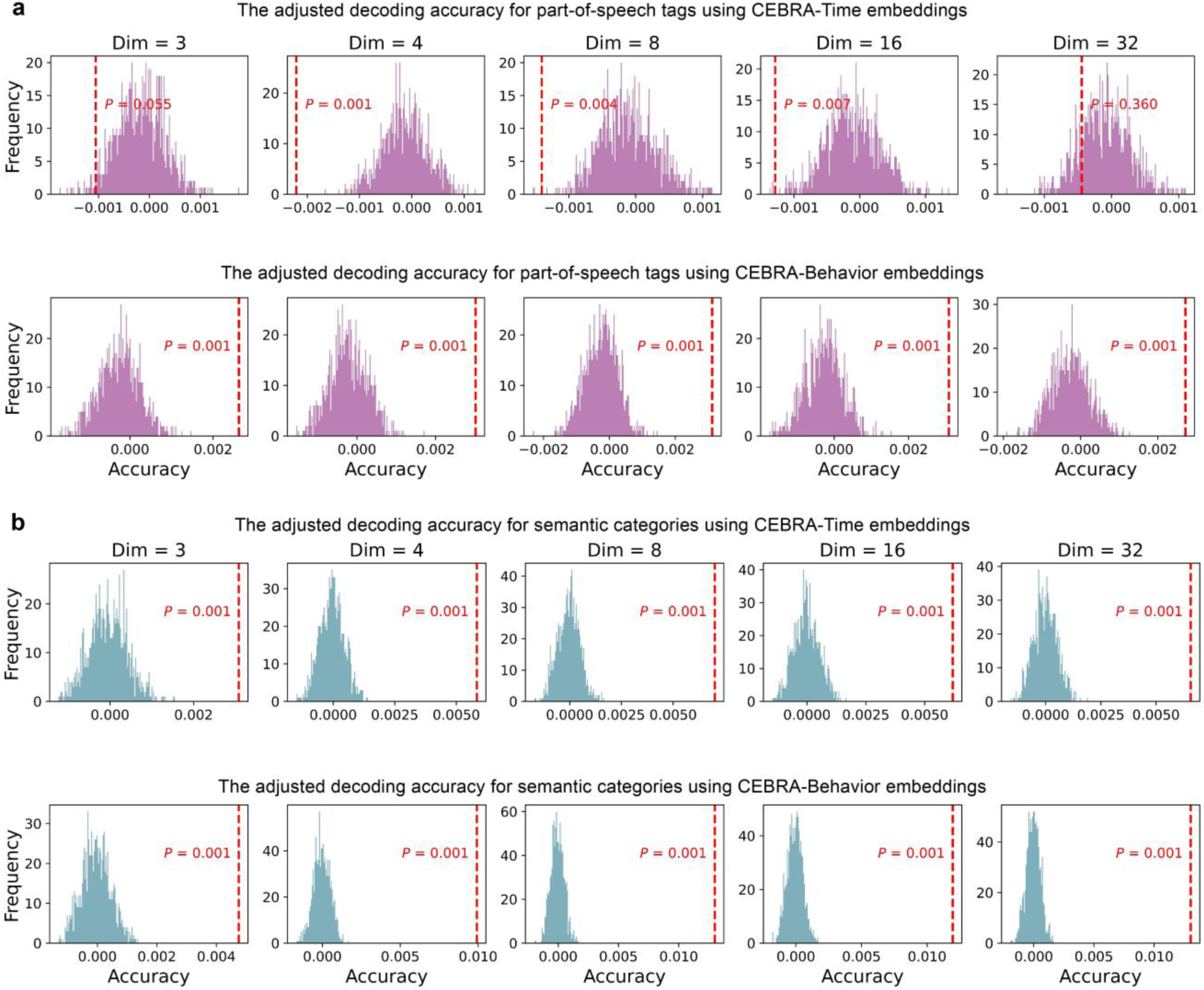
Adjusted decoding accuracies for two lexical features using CEBRA-Time and CEBRA-Behavior embeddings. **a)** Adjusted decoding accuracy for part-of-speech using CEBRA embeddings from 20 runs in the analysis dataset. **b)** Adjusted decoding accuracy for semantic classes. Two-tailed randomization tests were performed to determine whether the adjusted accuracy was significantly higher than chance. The null distributions were generated by randomly shuffling the target labels 1,000 times.

**Supplementary Figure 3.**
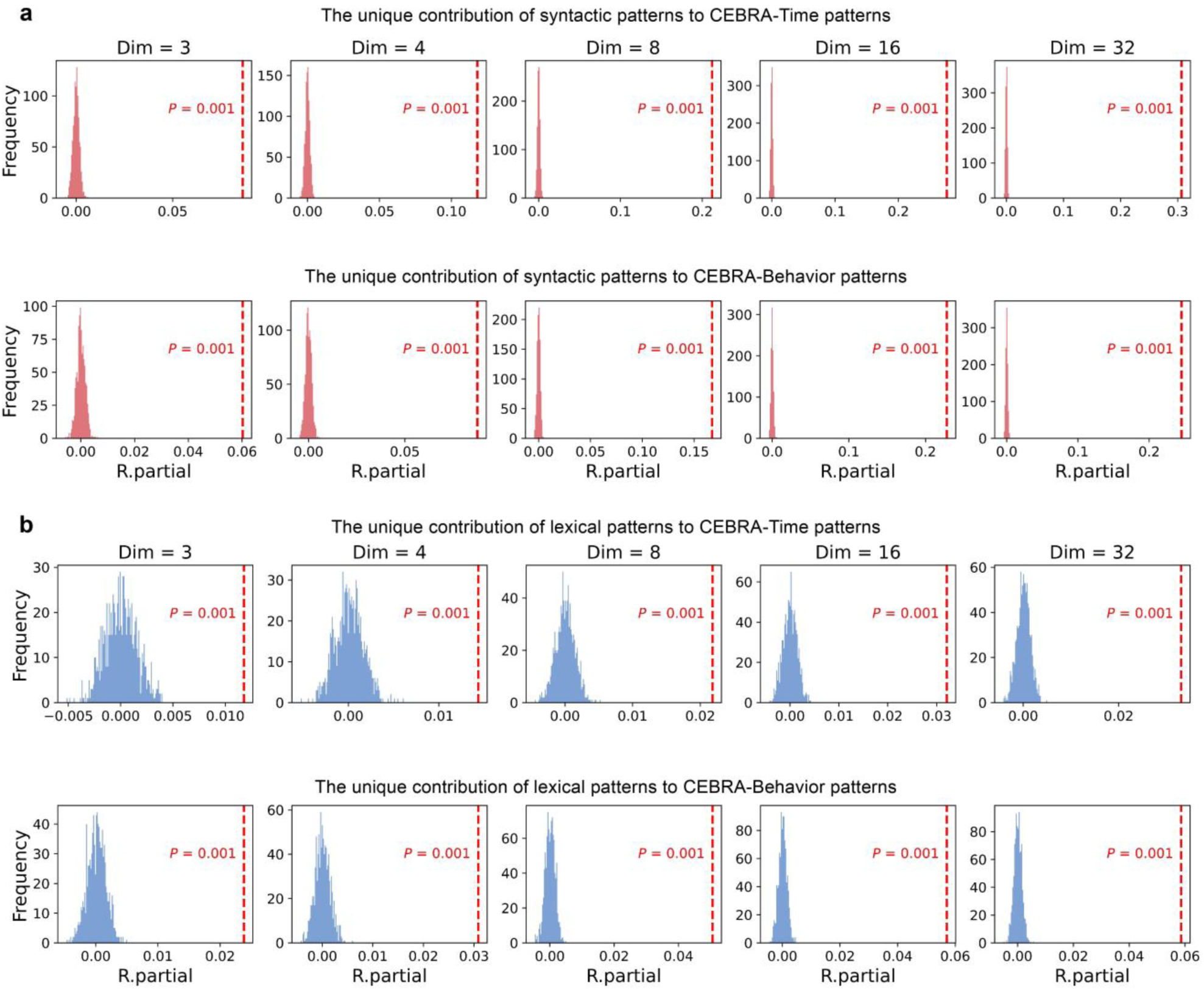
The semi-partial Spearman correlations between neural latent RDMs and two linguistic RDMs. **a)** Structured contributions to neural latent RDMs derived from CEBRA embeddings after removing the effects of acoustic feature, word length, word frequency, and distributed lexical representation. **b)** Distributed contributions to neural latent RDMs after removing the effects of acoustic feature, word length, word frequency, and syntactic relation. Two-tailed randomization tests were performed to determine whether the semi-partial correlation was significantly higher than chance. The null distributions were obtained by randomly shuffling each neural latent RDM 1,000 times before computing the semi-partial correlations.

**Supplementary Figure 4.**
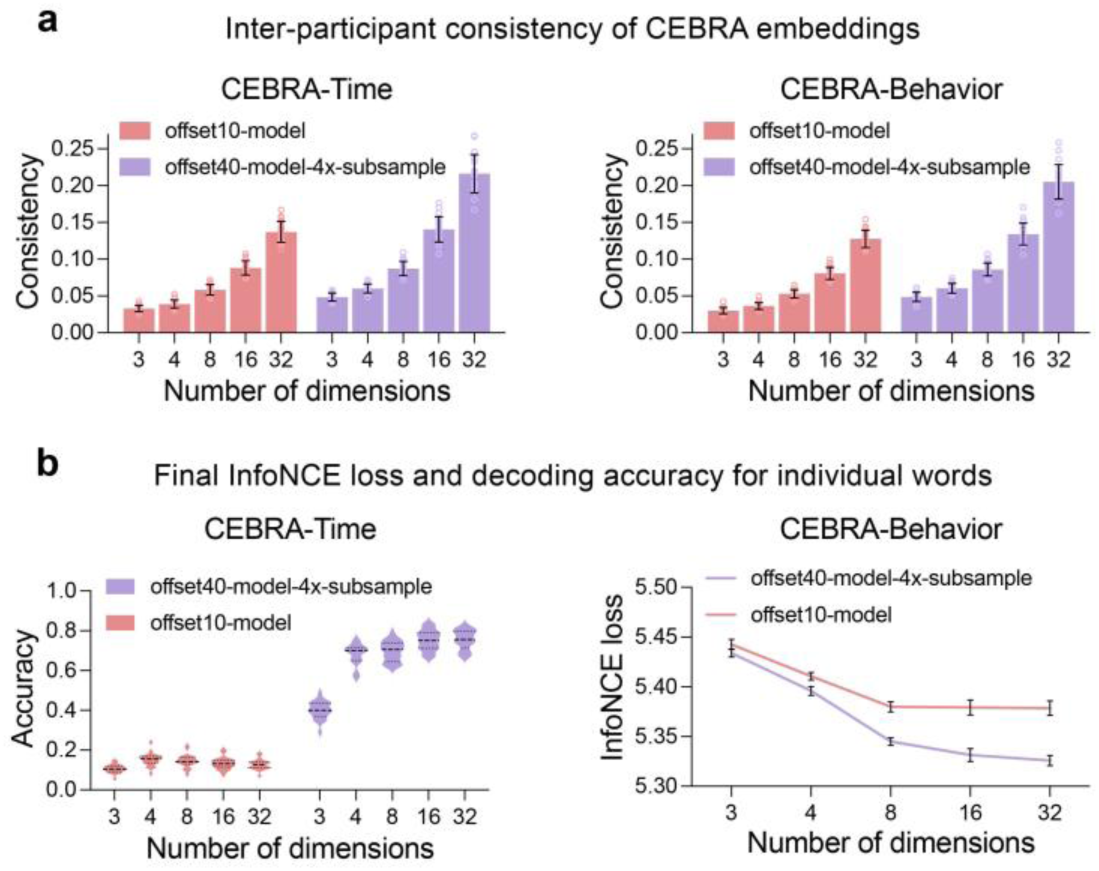
Exploration of language-relevant information in latent spaces derived from nonlinear encoders with different architectures. **a)** The inter-participant consistency of the CEBRA-Time and CEBRA-Behavior embeddings. The heights of the bars represent the average consistency values over 20 runs in the discovery dataset, with the error bars indicating the standard deviations. **b)** The word decoding accuracy using the CEBRA-Time embeddings and the final InfoNCE loss from behavioral contrastive learning, which were used to evaluate the ability of the CEBRA-Time and CEBRA-Behavior embeddings to capture language-relevant information. For the word decoding accuracy, the adjusted balanced accuracies within each run in the discovery dataset were averaged across participants to plot the violin plots. For the final InfoNCE loss, averages were also calculated within each run. The red and purple lines indicate the average loss values across 20 runs, and the black error bars indicate the standard deviations.

**Supplementary Figure 5.**
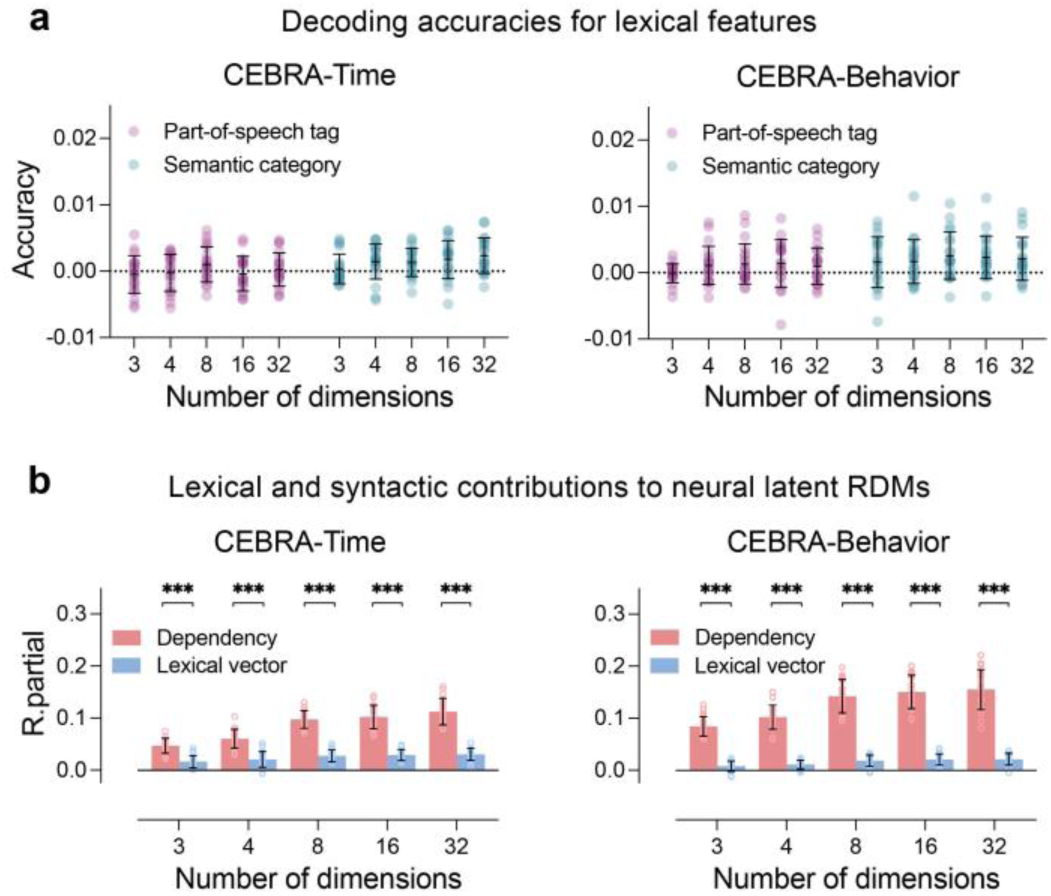
Decoding accuracies for lexical features and linguistic contributions to neural latent RDMs using an alternative setting (model: offset10-model, time offset = 10). **a)** Adjusted decoding accuracies for part-of-speech and semantic categories using word-level CEBRA embeddings from 20 runs in the analysis dataset. We observed significant but minor effect of attribute on decoding accuracies (*F*(1, 2360) = 16.458, *P* < 0.001, partial *η*^2^ = 0.007 for CEBRA-Time; *F*(1, 2360) = 8.840, *P* = 0.003, partial *η*^2^ = 0.004 for CEBRA-Behavior). The other effects were not significant (*Ps* > 0.05). Each observation represents the average decoding accuracy within each run, calculated across test sets and then across participants. The black lines represent the means and the standard deviations. **b)** Distributed and structured contributions to neural latent RDMs derived from CEBRA embeddings. We observed a significant interaction effect between linguistic feature and dimensionality (*F*(4, 2360) = 73.274, *P* < 0.001, partial *η*^2^ = 0.110 for CEBRA-Time; *F*(4, 2360) = 72.654, *P* < 0.001, partial *η*^2^ = 0.110 for CEBRA-Behavior). In addition, the structured syntactic RDMs explained significantly more variance than the distributed lexical RDMs in each dimension setting for both CEBRA-Time and CEBRA-Behavior (post-hoc Tukey’s HSD, *Ps* < 0.001). Furthermore, the syntactic contributions were more sensitive to the latent dimensionality (logarithmic model: standardized *β* = 0.553, SE = 0.023, *t*(1168) = 23.746, *P* < 0.001 for CEBRA-Time; *β* = 0.615, SE = 0.025, *t*(1168) = 25.062, *P* < 0.001 for CEBRA-Behavior) than lexical contributions (*β* = 0.246, SE = 0.030, *t*(1168) = 8.167, *P* < 0.001 for CEBRA-Time; *β* = 0.237, SE = 0.030, *t*(1168) = 8.012, *P* < 0.001 for CEBRA-Behavior). For visualization, the partial *R* values were averaged across participants within each run using data in the analysis dataset (*n* = 20).

**Supplementary Figure 6.**
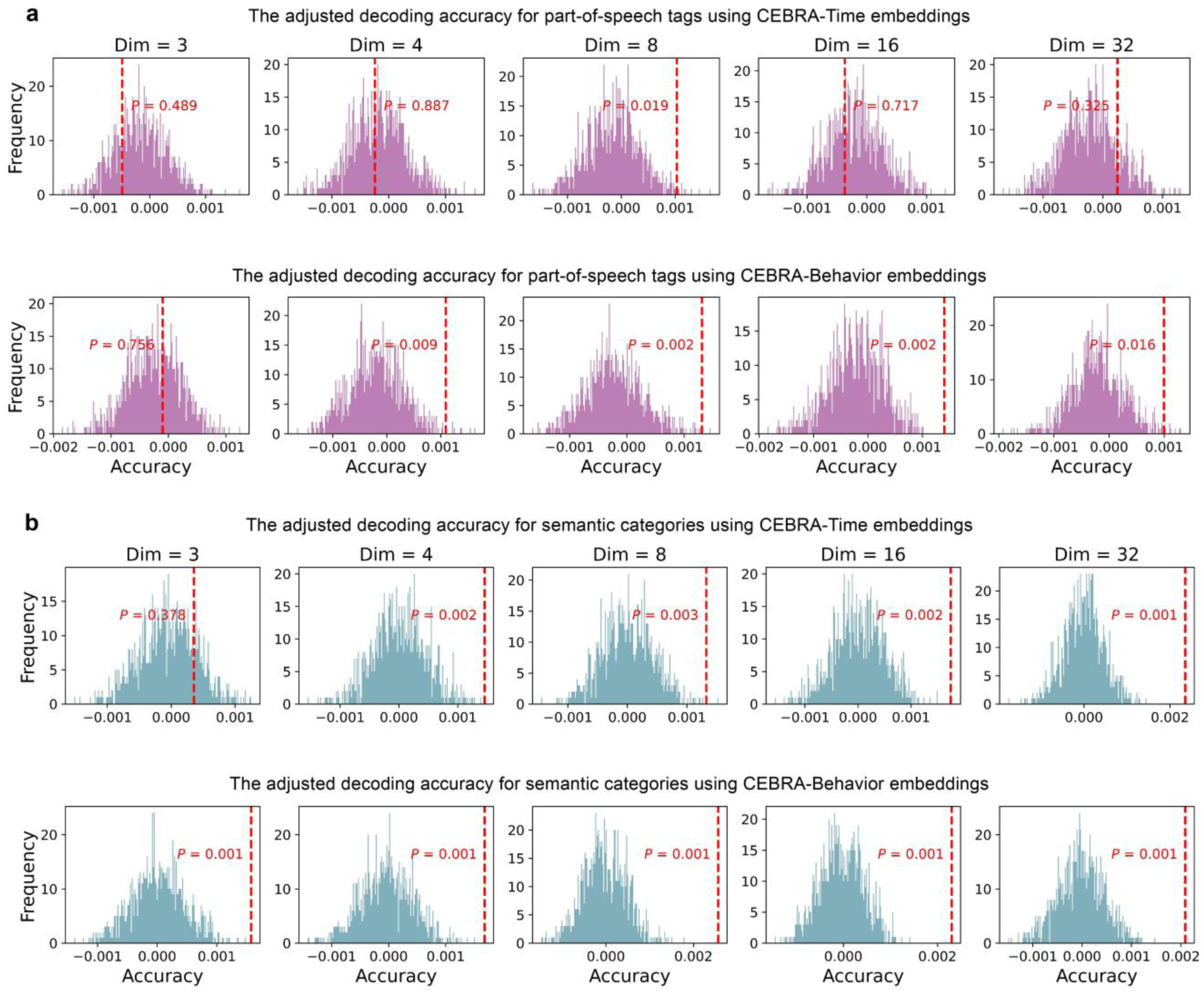
Adjusted decoding accuracies for two lexical features using CEBRA-Time and CEBRA-Behavior embeddings using an alternative setting (model: offset10-model, time offset = 10). **a)** Adjusted decoding accuracy for part-of-speech using CEBRA embeddings from 20 runs in the analysis dataset. **b)** Adjusted decoding accuracy for semantic classes. Two-tailed randomization tests were performed to determine whether the adjusted accuracy was significantly higher than chance. The null distributions were generated by randomly shuffling the target labels 1,000 times.

**Supplementary Figure 7.**
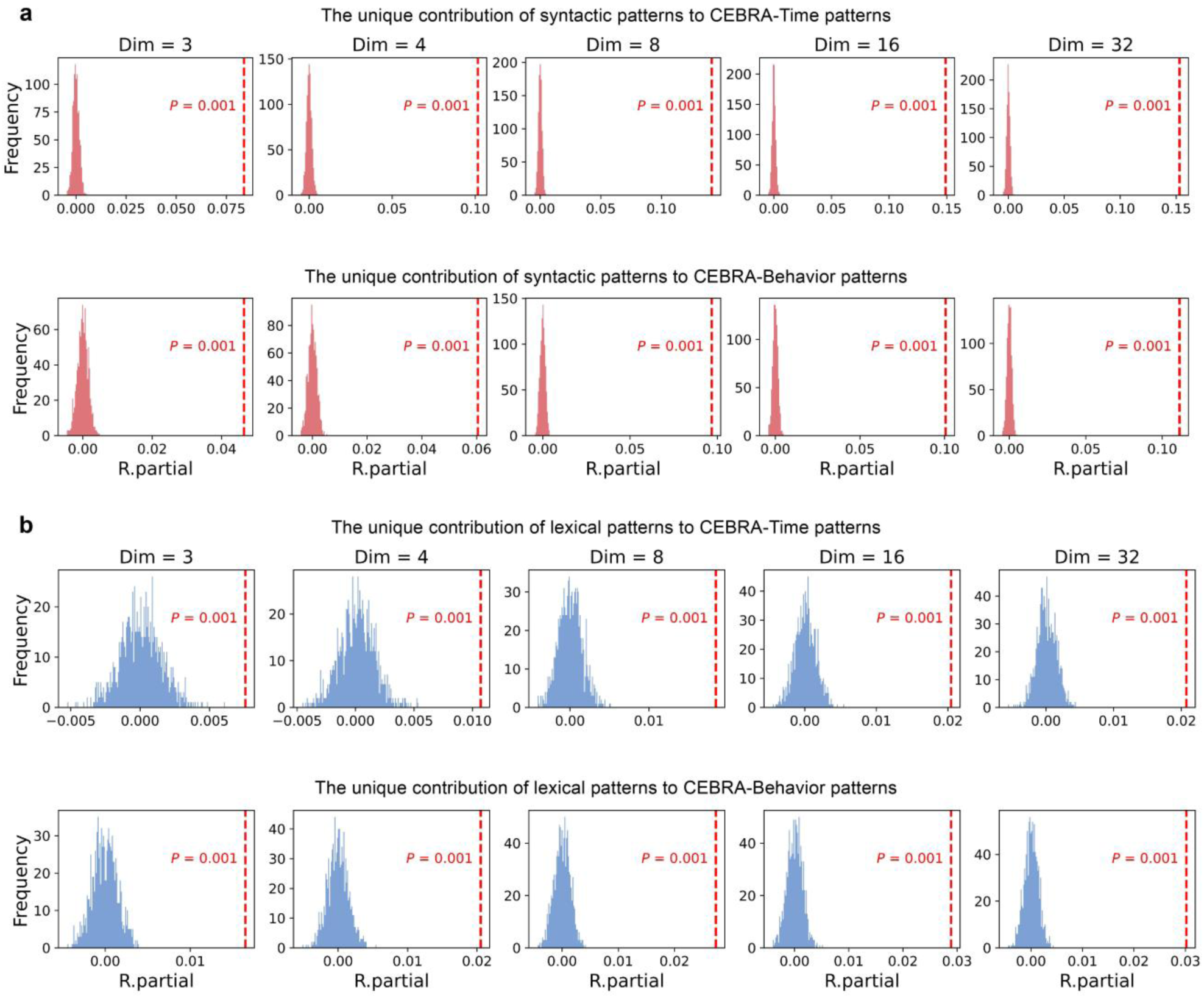
The semi-partial Spearman correlations between neural latent RDMs and two linguistic RDMs using an alternative setting (model: offset10-model, time offset: 10). **a)** Structured contributions to neural latent RDMs derived from CEBRA embeddings after removing the effects of acoustic feature, word length, word frequency, and distributed lexical representations. **b)** Distributed contributions to neural latent RDMs after removing the effects of acoustic feature, word length, word frequency, and syntactic relations. Two-tailed randomization tests were performed to investigate whether the semi-partial correlation was significantly higher than chance. The null distributions were obtained by randomly shuffling each neural latent RDM 1,000 times before computing the semi-partial correlations.

**Supplementary Figure 8.**
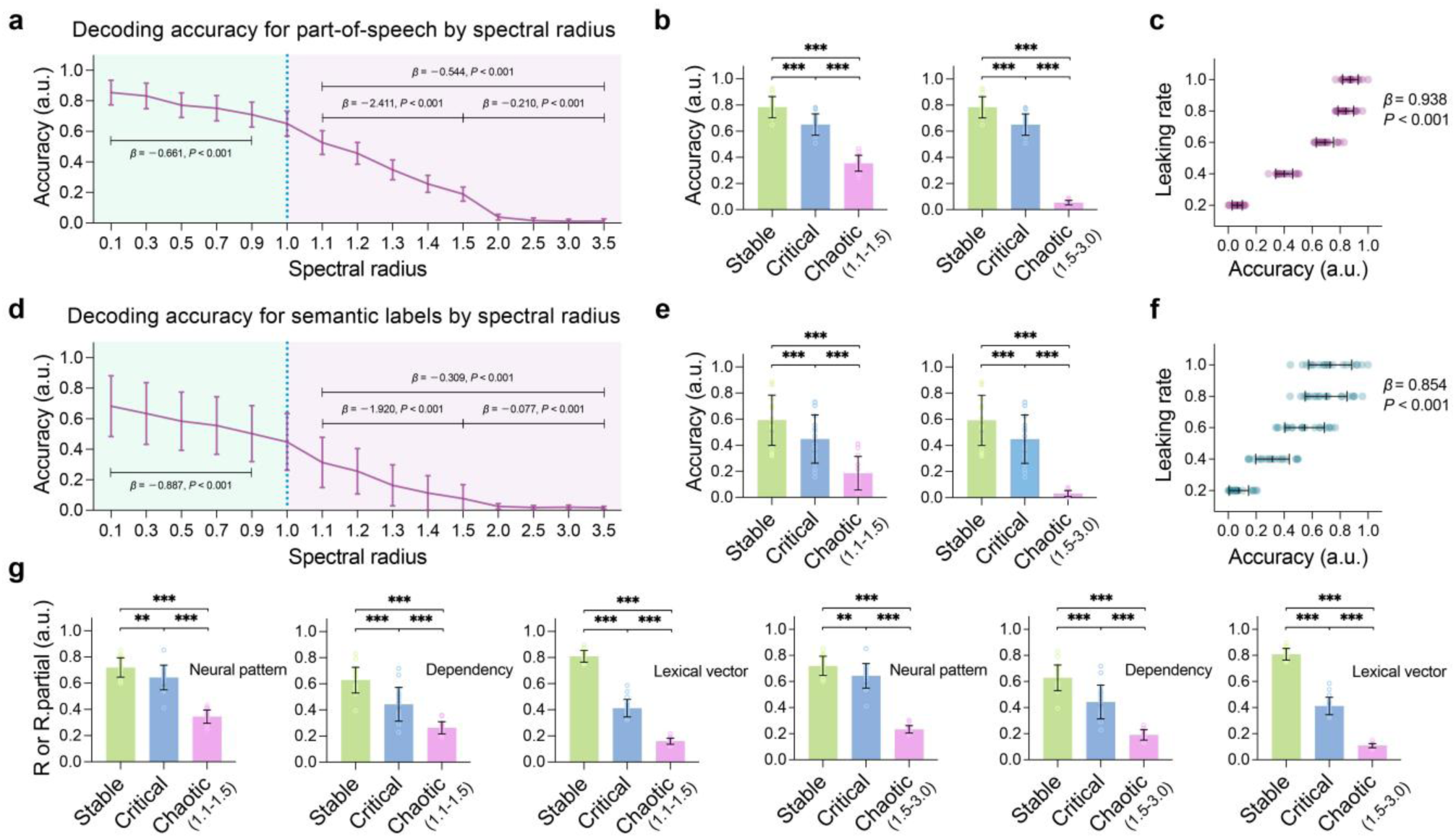
Decoding performance of lexical features using internal representations in brain-inspired RC systems by spectral radius and leaking rate. **a)** Adjusted decoding accuracies for part-of-speech as a function of the spectral radius with the leaking rate fixed at 0.4, using data from 20 runs in the simulation dataset, tested by linear mixed-effect models. The Fisher z-transformed (semi-partial) correlations were averaged for each run and then normalized across spectral radii for visualization. The lines and bars represent the means and the standard deviations. **b)** Pairwise comparisons between stratified states for the part-of-speech decoding accuracies using Wilcoxon signed-rank tests. The heights represent the average values over 20 runs, while the error bars indicate the standard deviations. Triple asterisks (***) indicate p-values less than 0.001, double asterisks (**) indicate p-values less than 0.01. **c)** Part-of-speech decoding across leaking rates, with the spectral radius fixed at 0.9, tested by linear mixed-effect models. Each observation represents the average score across participants within each run. The black lines represent the means and the standard deviations. **d-f)** Semantic decoding by spectral radius and leaking rate. **g)** Pairwise comparisons between stratified states for the brain-reservoir similarity, as well as the distributed and structured contributions to internal representations in brain-inspired RC systems.

**Supplementary Figure 9.**
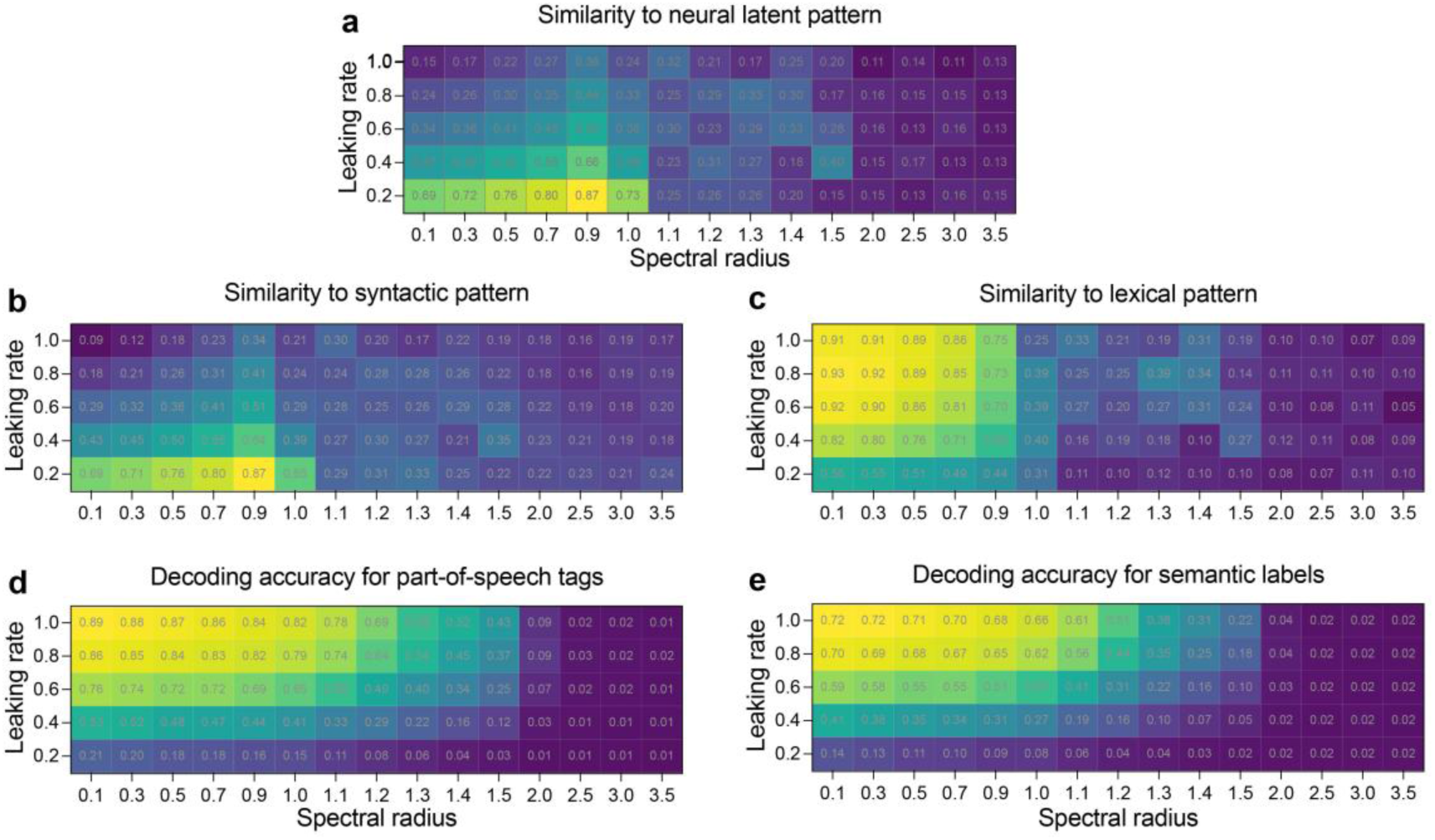
The effects of memory capacity and integration strategy on the internal representations in the brain-inspired RC systems. **a-e)** Heatmaps showing the brain-reservoir similarity, the unique contributions of distributed and structured RDMs, and decoding accuracies for part-of-speech and semantic categories, with their means plotted as a function of spectral radius (x-axis) and leaking rate (y-axis), using data from 20 runs in the simulation dataset. We observed significant interaction effects between leaking rate and stratified state on decoding accuracies for part-of-speech (*F*(2, 275) = 306.459, *P* < 0.001, partial *η*^2^ = 0.690) and semantic categories (*F*(2, 275) = 238.483, *P* < 0.001, partial *η*^2^ = 0.634). Within the stable state, the main effects of spectral radius and leaking rate on decoding accuracies were significant (*Ps* < 0.001, partial *η*^2^ ranging from 0.058 to 0.892), but the interaction effects were not significant (*F*(1, 477) = 0.676, *P* = 0.411, partial *η*^2^ = 0.001 for part-of-speech decoding; *F*(1, 477) = 2.609, *P* = 0.107, partial *η*^2^ = 0.005 for semantic decoding). For the five measures, the effect of leaking rate was significantly attenuated in the critical and chaotic states compared to the stable state (fixed effects in the linear mixed-effect model, *Ps* < 0.01), except for the effects of leaking rate on part-of-speech and semantic decoding when we configured the brain-inspired RC system to transition from the stable state to the critical state (*t*(275) = 0.160, *P* = 0.873 for part-of-speech decoding; *t*(275) = −0.001, *P* = 0.999 for semantic decoding).

**Supplementary Table 1.** The relationship between the dimensionality and information representations in neural latent spaces derived from nonlinear encoders with different architectures. The AIC and BIC were used to evaluate three candidate models characterizing mean-only, first-order polynomial, and logarithmic relationships. Mean-only models were defined as 𝑦 = 𝛽_0_ + 𝜀. First-order polynomial models were defined as 𝑦 = 𝛽_0_ + 𝛽_1_𝑥 + 𝜀. Logarithmic growth models were defined as 𝑦 = 𝛽_0_ + 𝛽_1_ln (𝑥) + 𝜀. The values represent the AIC/BIC metrics corresponding to each model, with those in bold font indicating the model with the best fit. The primary results from the offset40-model-4x-subsample architecture are presented in the main text. The results from the offset10-model architecture are reported here, which largely confirm main findings, although it performs less well in capturing consistent and language-related latent components of neocortical responses.

| Encoder | CEBRA-Time (offset40-model-4x-subsample) |  |  |  |  |  |
| --- | --- | --- | --- | --- | --- | --- |
| Measures | Consistency | Individual word | Part-of-speech | Semantic classes | Lexical content | Syntactic relations |
| Mean-only | 3411.236 / 3431.597 | 3265.070 / 3285.430 | **3356.197 / 3376.558** | 3259.314 / **3279.674** | 3243.601 / 3263.962 | 3368.550 / 3388.911 |
| Linear | **472.401 / 497.851** | 3054.167 / 3079.617 | 3363.888 / 3389.339 | 3266.479 / 3291.930 | 3123.027 / 3148.477 | 2200.697 / 2226.147 |
| Logarithmic | 855.064 / 880.514 | **2903.457 / 2928.908** | 3360.045 / 3385.495 | **3257.340 / 3282.791** | **3082.722 / 3108.172** | **1557.711 / 1583.161** |
| Encoder | CEBRA-Time (offset10-model) |  |  |  |  |  |
| Mean-only | 3408.523 / 3428.883 | **2871.225 / 2891.586** | **3392.465 / 3412.825** | **3411.348 / 3426.618** | 3313.082 / 3333.442 | 3127.639 / 3147.999 |
| Linear | **401.108 / 426.558** | 2882.963 / 2908.414 | 3404.328 / 3429.779 | 3418.238 / 3438.598 | 3279.946 / 3305.397 | 2836.714 / 2862.165 |
| Logarithmic | 833.857 / 859.307 | 2877.500 / 2902.951 | 3399.035 / 3424.486 | 3412.684 / 3433.044 | **3255.340 / 3280.790** | **2674.839 / 2700.290** |
| Encoder | CEBRA-Behavior (offset40-model-4x-subsample) |  |  |  |  |  |
| Measures | Consistency | InfoNCE Loss | Part-of-speech | Semantic classes | Lexical content | Syntactic relations |
| Mean-only | 3406.643 / 3427.004 | 3408.452 / 3418.632 | **3357.350 / 3372.620** | 3221.110 / 3241.471 | 3131.149 / 3151.509 | 3379.390 / 3399.750 |
| Linear | **469.845 / 495.295** | 2464.697 / 2479.968 | 3369.398 / 3389.758 | 3204.587 / 3230.038 | 2904.342 / 2929.793 | 2242.682 / 2268.133 |
| Logarithmic | 837.875 / 863.326 | **1553.467 / 1568.737** | 3364.405 / 3384.765 | **3182.409 / 3207.859** | **2788.713 / 2814.163** | **1561.837 / 1587.287** |
| Encoder | CEBRA-Behavior (offset10-model) |  |  |  |  |  |
| Mean-only | 3414.305 / 3434.666 | 3413.537 / 3433.897 | **3402.803 / 3423.163** | **3380.273 / 3400.634** | 3278.704 / 3299.064 | 3280.530 / 3300.891 |
| Linear | **358.842 / 384.292** | 2921.373 / 2946.823 | 3414.350 / 3439.800 | 3392.074 / 3417.524 | 3246.737 / 3272.187 | 2940.614 / 2966.064 |
| Logarithmic | 956.955 / 982.406 | **2447.545 / 2472.995** | 3408.454 / 3433.904 | 3386.594 / 3412.045 | **3223.365 / 3248.815** | **2785.058 / 2810.508** |

